# Tauopathy primes co-filament assembly and dysfunction of TDP-43

**DOI:** 10.64898/2026.05.11.723888

**Authors:** Meghraj S. Baghel, Grace D. Burns, Aswathy Peethambaran Mallika, Xiaoke K. Chen, Fangbing Liu, Shruti Renganathan, Tong Li, Juan C. Troncoso, Philip C. Wong

## Abstract

While most Alzheimer’s disease (AD) which is associated with <u>L</u>imbic <u>P</u>redominant <u>A</u>ge-related <u>T</u>DP-43 <u>E</u>ncephalopathy (LATE) exhibits accelerated brain atrophy, the pathogenic mechanism remains elusive. We show here, in mice harboring depositions of amyloid-β and tau, the age-dependent emergence of TDP-43 proteinopathy. We demonstrate that TDP-43 dysfunction facilitates caspase 3-mediated endoproteolysis of tau, accelerates tauopathy and exacerbates neuron loss. Unexpectedly, we found that the emergence and spread of TDP-43 proteinopathy is associated with the spread of tauopathy and correlated with co-filament assembly of tau and TDP-43. Importantly, TDP-43 dysfunction precedes such co-filament assembly and TDP-43 cytoplasmic aggregates. Consistent with the idea that tauopathy could prime co-filament assembly and proteinopathy of TDP-43 to exacerbate neurodegeneration, we found tau co-filament assembly with TDP-43 in AD and AD-LATE cases. These findings suggest that TDP-43 dysfunction accelerates tauopathy, which, in turn, primes co-filament assembly and dysfunction of TDP-43 to exacerbate neuron loss in AD-LATE, a pathogenic mechanism disclosing novel targets and therapeutic strategies.

## Introduction

Cytoplasmic aggregation of TDP-43 (*1*), an RNA/DNA binding protein that regulates the transcriptome by repressing non-conserved cryptic exons (*2*), is found in numerous neurodegenerative disorders. Genetic, biomarker, and neuropathological data indicate that loss of TDP-43 splicing repression begins before symptoms and contributes to neuron loss in ALS, FTLD-TDP, and Alzheimer’s disease (AD)-LATE (*3–16*). Up to 75% of AD and AD-related dementia (ADRD) cases show non-canonical pathologies, including TDP-43 in LATE (*17*), α-synuclein Lewy bodies, and tau pick bodies (*18–22*). TDP-43 pathology (cytoplasmic aggregation) occurs in 40–60% of ADRD cases and correlates with worsened neurodegeneration and cognitive decline (*18–20*). Such TDP-43 proteinopathy is also reported in other tauopathies such as frontotemporal lobar dementia (FTLD), corticobasal degeneration (CBD), primary age-related tauopathy (PART), and progressive supranuclear palsy (PSP) (*23–26*). Previously, we found deposition of Aβ plaque facilitates tau pathology (*27–28*), and recent studies demonstrated that loss of TDP-43 promoted the pathological tau conversion to exacerbate loss of vulnerable neurons in mouse models (*29*), suggesting a mechanism to account for the accelerated tauopathy in driving neuron loss in AD-LATE. However, a critical unresolved question is how accelerated tauopathy would lead to the development of TDP-43 proteinopathy in AD-LATE.

Because TDP-43 dysfunction occurs in some neurons of the human aging brain (*6*), we asked whether tauopathy could lead to TDP-43 cytoplasmic aggregation during aging. We addressed this question using our previously established mouse model exhibiting amyloidosis and tauopathy and found in these aged mice the emergence of TDP-43 cytoplasmic aggregates in vulnerable neurons that is dependent on tauopathy. Importantly, we provide evidence of co-filament assembly of tau and TDP-43 occurring in mouse models as well as in AD brains, suggesting a potential pathogenic mechanism to drive TDP-43 cytoplasmic aggregation and dysfunction. We thus propose a feed forward mechanism whereby TDP-43 dysfunction accelerates tauopathy, which, in turn, drives TDP-43 proteinopathy to exacerbate neuron loss in AD-LATE.

## Results

### TDP-43 dysfunction and cytoplasmic aggregation occur in aged *Tau4R;AP* mice

While recent studies suggested that tau aggregates may provoke TDP-43 proteinopathy to exacerbate neuron loss (*32–33*) in AD-LATE cases (*18–20*), how the co-pathology of TDP-43 confers such a negative impact remains poorly understood. To address this question, we first ask whether aging could be a contributing factor in AD-LATE since we previously documented that TDP-43 dysfunction occurs in the human aging brain (*6*). Taking advantage of an *in vivo* tau seeding model expressing low level of a human tau 4-repeat fragment (*Tau4R* mice) when challenge by Aβ plaque (*Tau4R-AP* mice) facilitates pathological conversion of endogenous tau (*27*), we asked whether the emergence of TDP-43 proteinopathy is associated with tauopathy in these aged *Tau4R;AP* mice. Using a well-established antisera recognizing misfolded TDP-43 at pS409/410, while small, immature neuronal cytoplasmic aggregates were detected in 20 month-old *Tau4R;AP* mice, more mature cytoplasmic aggregates can be readily observed in different brain regions including hippocampus and cortex by 24 months of age (**Fig. 1a**, **second and third column**); however, no such immunoreactivities were observed in age matched wild-type (WT) littermates (**Fig. 1a**, **first column**). Notably, the different patterns of the pTDP-43 aggregates in neurons of aged *Tau4R-AP* mice were observed which are reminiscent of those found in human brains of FTLD or AD cases (*1, 18, 34–39*) such as small pTDP-43 cytoplasmic inclusions (**Fig. 1b, panel I**) and cytoplasmic aggregates with and without skein-like morphology within axon (**Fig 1b, panels II & III; Sup Fig. 1b**) which resemble an advanced stage of aggregation. Thus, we surmise that these small pTDP-43 cytoplasmic inclusions (**Fig. 1a; Sup Fig. 1b; Sup Fig. 2a, c-d; first column**) evolves to mature aggregates when these mice age.

**Figure 1.**
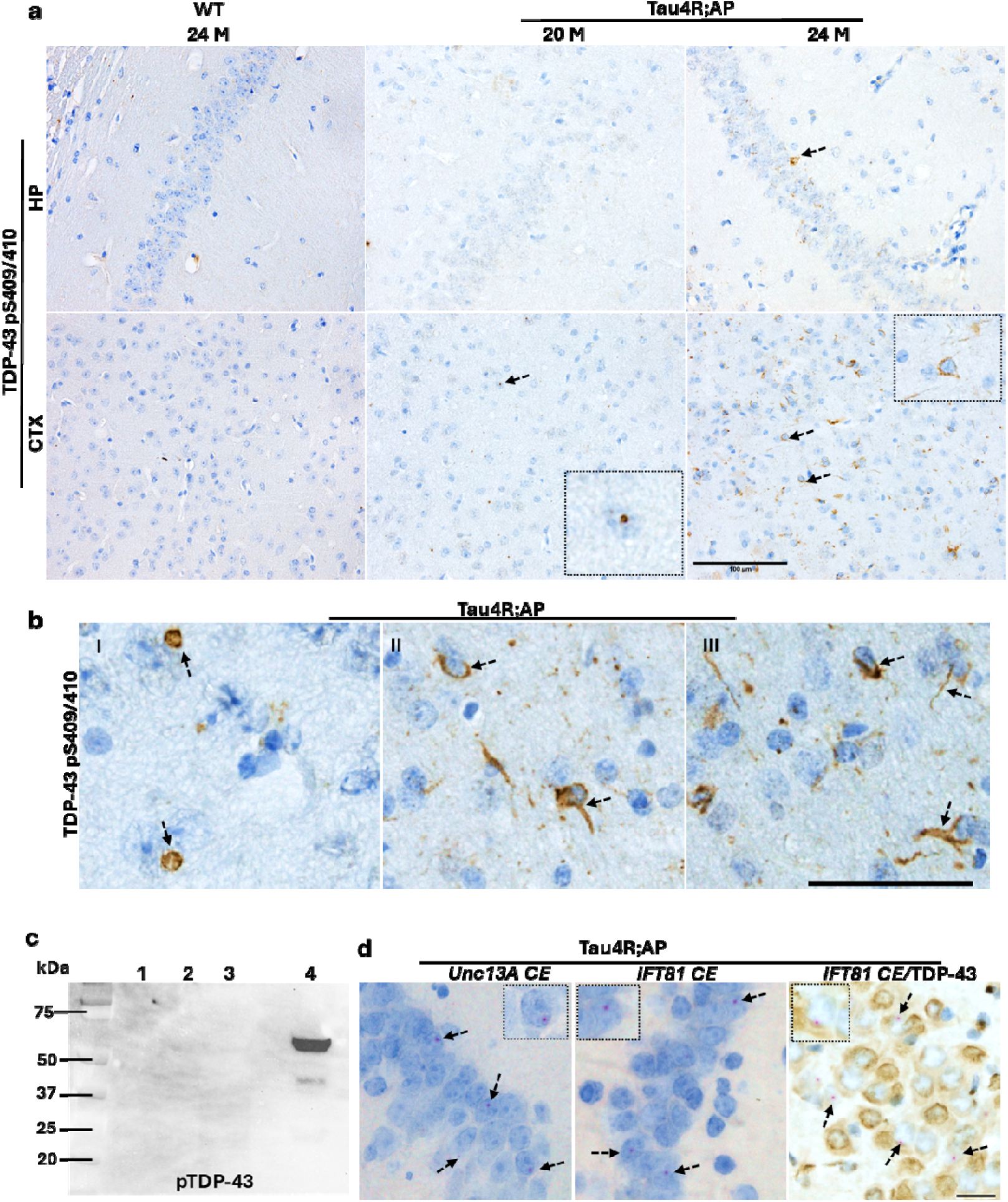
TDP-43 dysfunction and cytoplasmic aggregation occur in aged *Tau4R;AP* mice. **(a)** Immunohistochemical analysis of brains of 20 and 24-month-old *WT* (n=5), *Tau4R;AP* (n=3) mice using antibody specific to pTDP-43 409/410 to detect cytoplasmic aggregation in the hippocampus and cortex (Arrow heads show the pTDP-43 cytoplasmic aggregations; scale bar, 100μm). **(b)** Immunohistochemistry of pTDP-43 showing different types of patterns of aggregations (Arrow heads indicate cytoplasmic inclusion; scale bar, 50μm). (**c)** Immunoblotting of Sarkosyl-insoluble extraction of pTDP-43 aggregates from Tau4R;AP and respective controls. Loading in Lane 1: WT, P3; 2: TDP-43 cKO, P3; 3: Tau4R;AP, S3; 4: Tau4R;AP, P3 of 24-month-old. (**d**) RNA-In situ hybridization analysis of brain sections from Tau4R:AP 24-month-old mice showing the presence of *Unc13a* and *Ift81* cryptic exons and *Ift81* CE and TDP-43 protein co-detection in the brain. (Arrow heads indicate pink signal of cryptic exon RNA inclusions; scale bar, 20μm)

**Figure 2:**
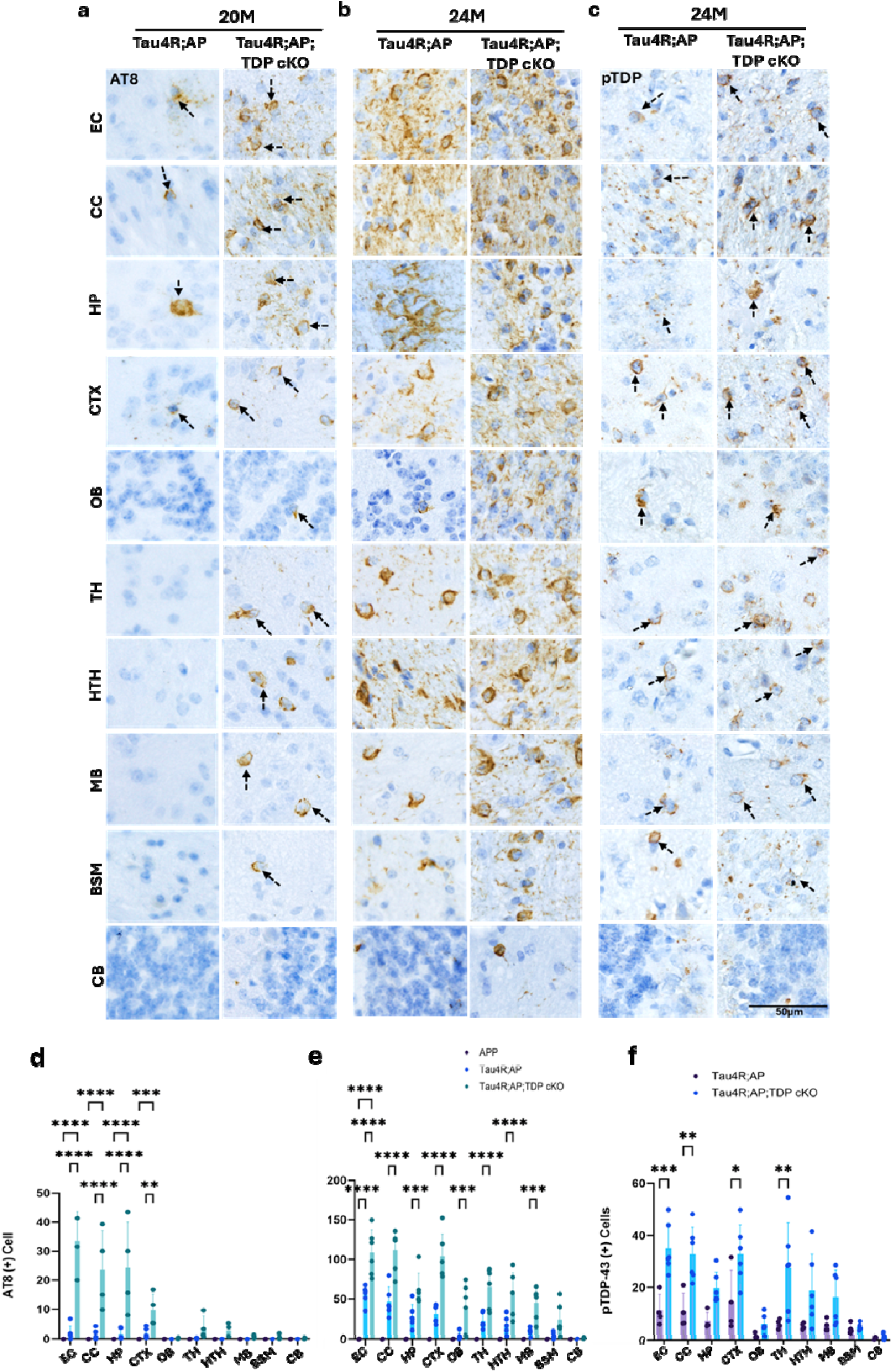
Emergence and spread of TDP-43 proteinopathy are dependent on the spread of tauopathy. **(a-b)** Immunohistochemical analysis showed age-dependent spreading of endogenous tau tangle in Tau4R-AP and Tau4R-AP;TDP cKO 20, 24-months old mice using antibodies specific to endogenous phosphorylated tau (AT8). The brain regions are entorhinal cortex (EC); corpus callosum (CC); hippocampus (HP); cerebral cortex (CTX); olfactory bulb (OB); thalamus (TH); hypothalamus (HTH); midbrain (MB); brain stem (BSM), and cerebellum (CB). Tau tangles first appeared in EC, CC, CTX and HP regions of Tau4R;AP, which was accelerated in further brain regions including OB, TH, HTH including MB and BSM in Tau4R;AP;TDP cKO mice (20 months of age, first two columns). With aging, tau tangles spread were extended in brain region OB, TH, HTH including MB and BSM in even Tau4R;AP 24-month-old mice, which was accelerated in Tau4R;AP;TDP cKO mice (middle two columns). (**c**) Immunohistochemical analysis showed age-dependent spreading of endogenous tau tangle in Tau4R-AP and Tau4R-AP;TDP cKO 20, 24-months old mice using antibodies specific to pTDP-43 409/410. The immunoreactivity of mature aggregated pTDP-43 was observed in brain regions EC, CC, HP, CTX), OB, TH, HTH, MB and BSM in Tau4R;AP 24-month-old, which was accelerated in Tau4R;AP;TDP cKO mice (last two columns). Arrow heads indicate the immunoreactivity of AT8 and pTDP-43, scale bar, 50um. (**d-e**) Quantification of AT8 positive cells in different brain regions of Tau4R;AP and Tau4R;AP;TDP cKO mice 20-, and 24-month-old respectively (two-way ANOVA; ns: no significant difference; *P<0.05; ****P<0.0001). (**f**) Quantification of pTDP-43 aggregate positive cells in different brain regions of Tau4R;AP and Tau4R;AP;TDP cKO 24-month-old mice. (two-way ANOVA; ns: no significant difference; *P<0.05; ****P<0.0001).

To confirm that these pTDP-43 aggregates are sarkosyl-insoluble, we performed differential protein solubility studies using brain extracts from *Tau4R;AP* mice. As expected, we detected pTDP-43 only in the sarkosyl-insoluble fraction (P3) of *Tau4R;AP,* but not WT or TDP-43cKO, mice (**Fig. 1c**). Since TDP-43 aggregates is accompanied by nuclear loss of TDP-43 function as indicated by inclusion of cryptic exons (*2–11*), we tested whether TDP-43 dysfunction occurs in neurons of *Tau4R;AP* mice. We assessed the inclusion of TDP-43 dependent cryptic exons we previously documented to occur within the mouse *Unc13a* or *Ift81* pre-mRNA (*40*) by RNA i*n-situ* hybridization (BaseScope method). The inclusion of *Unc13a* or *Ift81* cryptic exon accompanied by nuclear clearance of TDP-43 was observed in the brain of *Tau4R;AP* mice (**Fig. 1d**). These observations suggest that canonical amyloid-β and tau pathologies precede TDP-43 proteinopathy in an age-dependent manner offering the possibility that in AD-LATE, tauopathy facilitates the emergence of TDP-43 proteinopathy.

### Tauopathy is sufficient to drive TDP-43 proteinopathy

To test whether TDP-43 proteinopathy is dependent on tauopathy, we assessed the emergence of TDP-43 cytoplasmic aggregation in *Tau4R;AP* mice relative to the onset of Aβ or tau pathology. As previously observed in *APP;PS1* or *Tau4R;AP* mice, Aβ deposition emerged ∼ 6-months of age (*27*); however, by 20 months we still failed to detect mature pTDP-43 aggregates in neurons of *Tau4R;AP* mice. In contrast, tauopathy emerged by 20 months (**Sup Fig. 3a, first column**) which spread in an age-dependent manner to different brain regions and by 24 months reaching the brain stem (**Sup Fig. 3a, second column; Sup Fig. 4a-b, first column**). Notably, during early stage of tau pathology, small cytoplasmic pTDP-43 inclusions can be observed (**Sup Fig. 3b, first column**) and as tauopathy advanced, these small inclusions seemed to be evolved into more mature cytoplasmic aggregates by 24 –months (**Sup Fig. 3b, second column**). Furthermore, we observed pTDP-43 immunoreactivity colocalized with tau aggregates (AT8) as tauopathy spreads from the cortex and hippocampus to the brain stem of *Tau4R;AP* mice (**Sup Fig. 3c**) suggesting that the tau pathology primes TDP-43 proteinopathy.

**Figure 3:**
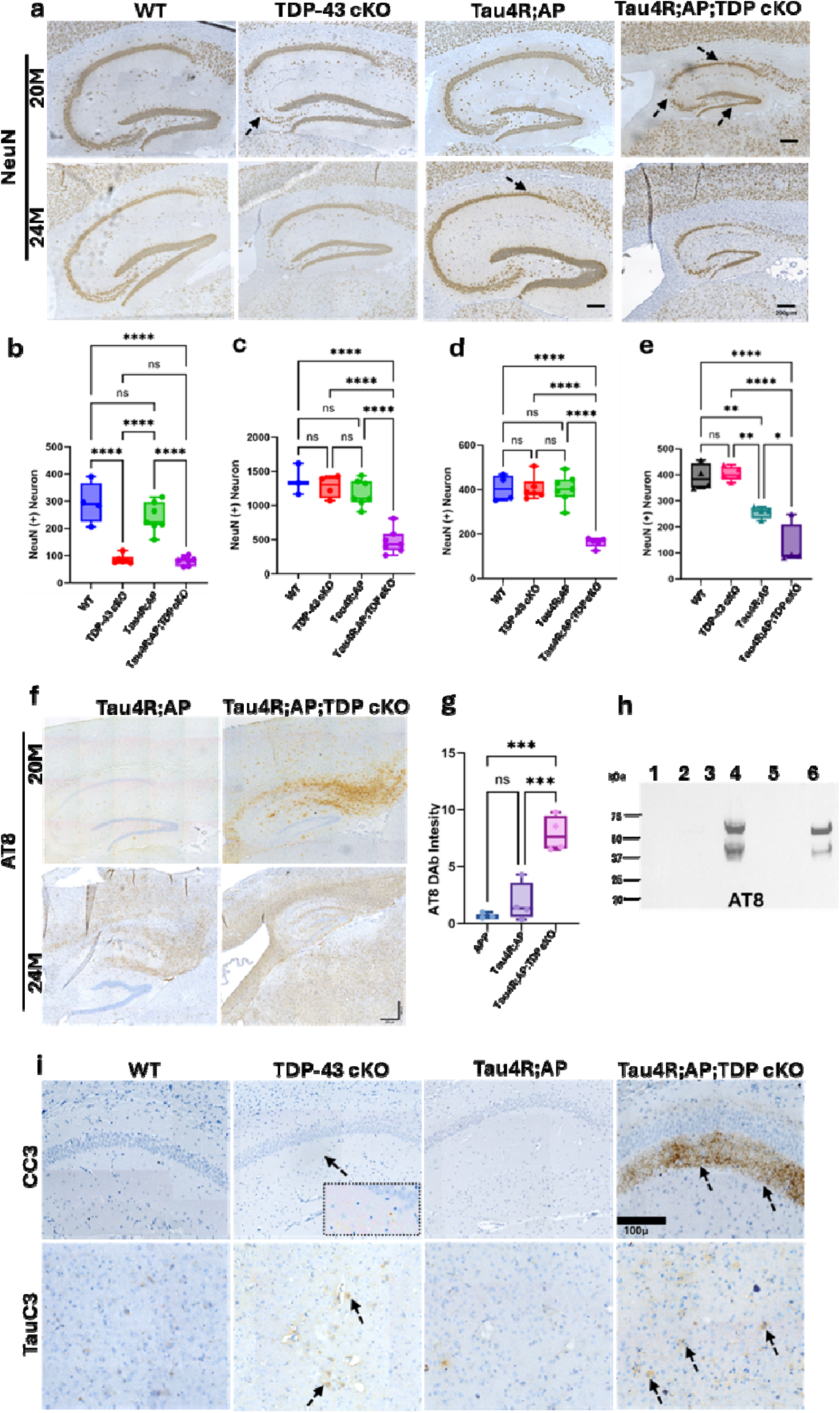
TDP-43 dysfunction accelerates tauopathy and TDP-43 proteinopathy to exacerbate neuron loss. **(a)** Immunohistochemical analysis of brains of 20 and 24-month-old *WT* (n=4), TDP-43 cKO (n=6), Tau4R;AP (n=6) and *Tau4R;AP;TDP cKO* mice (n=6) using antibody specific to NeuN to detect neurons (Arrow heads indicates neuronal loss in hippocampal subfields, Scale bar, 200 μm). **(b-d)** Quantification of NeuN positive neurons from CA2/3, DG and CA1 respectively in 20-month-old mice, **(e)** NeuN positive neuron quantification hippocampal CA1 in 24-month-old mice. (one-way ANOVA; ns: no significant difference; *P<0.05; ****P<0.0001). (**f)** Immunohistochemical analysis of the hippocampus using antibodies specific to endogenous phosphorylated tau, AT8 (upper panel, 20-months) and (lower panel, 24-months) in Tau4R;AP and Tau4R;AP;TDP cKO (Scale bar, 200 μm). (g) Quantification of AT8 immunoreactivity in 20-month-old APP, Tau4R;AP and Tau4R;AP;TDP cKO (one-way ANOVA; ns: no significant difference considered at P<0.05). **(h)** Immunoblotting of Sarkosyl-insoluble extraction of pTau aggregates from Tau4R;AP, Tau4R;AP;TDP cKO and respective controls. Loading in lane 1: WT, P3; 2: TDP-43 cKO, P3; 3: Tau4R;AP, S3; 4: Tau4R;AP, P3; 5: Tau4R;AP;TDP cKO, S3; 6: Tau4R;AP;TDP cKO, P3 of 24-month-old. (**i**) Immunohistochemical analysis using cleaved caspase 3 (CC3) brain sections of WT, TDP-43 cKO, Tau4R;AP and Tau4R;AP;TDP cKO 20-month-old mice (upper panel). The lower panel shows the immunoreactivity of cleaved tau (TauC3). Arrow heads indicate the immunoreactivity signal of CC3 and TauC3, scale bar, 100um.

**Figure 4:**
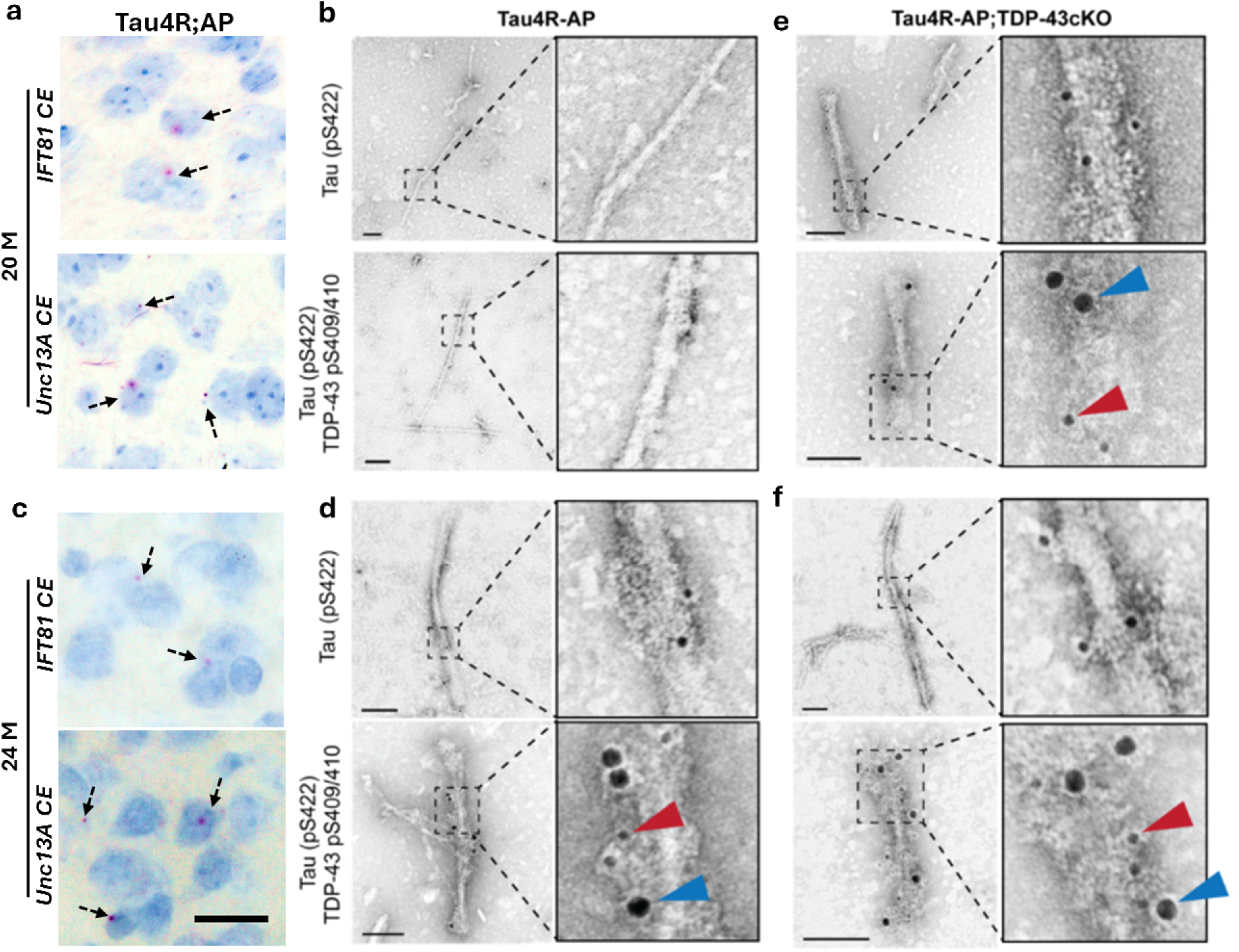
Tauopathy primes dysfunction and co-filament assembly of TDP-43. BaseScope analysis of cryptic RNA detection of *Ift81 and Unc13a* in the brain of **a)** 20- and **c)** 24-month old Tau4R-AP mice (arrow heads indicate the pink RNA foci, scale bar, 20µm). Double labeling immuno-EM analysis of sarkosyl-insoluble brain extracts derived from **b, e)** 20-month-old and **d, f)** 24-month-old Tau4R-AP and Tau4R-AP;TDP-43cKO mice using primary antibodies against tau (pS422) and TDP-43 (pS409/410) and goat anti-rabbit 6nm gold-conjugated (red arrowheads) and goat anti-mouse 12nm gold-conjugated (blue arrowheads) secondary antibodies, respectively (scale bar, 100nm).

To test directly whether tauopathy is sufficient to drive pTDP-43 proteinopathy, we took advantage of our previous AAV-hTau4R induced tau seeding model (*29*) where AAV mediated expression of hTau4R facilitated pathological conversion of endogenous tau in brains of wild-type mice. Such AAV-PHP.eB-hTau4R model facilitated robust pathological conversion of endogenous tau (10 months post injection; mpi) observed in different brain regions, including the hippocampus (**Sup Fig. 3d, upper panel**). Importantly, we observed emergence of pTDP-43 cytoplasmic aggregates that was dependent on tauopathy in neurons of this tauopathy mouse model (**Sup Fig. 3d, lower panel**), suggesting that tauopathy, independent of amyloid-β plaque, is sufficient to prime the cytoplasmic pTDP-43 aggregates to exacerbate neuron loss in AD-LATE.

### The emergence and spread of TDP-43 proteinopathy are dependent on the spread of tauopathy

To determine whether cytoplasmic pTDP-43 pathology is dependent on tau pathology in *Tau4R-AP* mice, we examined initially the relationship between pTDP-43 immunoreactivity and tau aggregation. In 20 months old *Tau4R;AP* mice, small TDP-43 inclusions were found corelated with tau aggregates in different brain regions (**Fig. 2a, first column; Sup Fig. 2a, c, first columns**). By 24 months, small inclusions of TDP-43 seem to evolve into mature cytoplasmic aggregates, the emergence of which is dependent on the spreading of tau pathology in different brain regions of 24–month old *Tau4R;AP* mice (**Fig. 2b-c, first column; Sup Fig. 2b, d, first column & 4b**).

If the emergence of TDP-43 proteinopathy indeed depends on tauopathy, we asked whether by accelerating the spread of tauopathy in these *Tau4R;AP* mice, the spread of TDP-43 proteinopathy would follow the spread of tauopathy. To test this prediction, we generated *Tau4R;AP* mice lacking TDP-43 in their forebrain neurons (*Tau4R;AP;CamKII-CreER;Tardbp^f/f^* or *Tau4R;AP;TDP43cKO*) and control littermates (**Sup Fig. 1a**). We depleted TDP-43 at 12-month of age before tau pathology initiates in *Tau4R;AP* mice (**Sup Fig. 3a**) and analyzed tau and TDP-43 pathologies at 20- and 24-months of age. While mature pTau aggregates were restricted to a paucity of cells in entorhinal cortex (EC), corpus callosum (CC), hippocampus (HP) and cortex (CTX) of 20-months old *Tau4R;AP* mice, marked increases in tau burden are found not only in these same regions of *Tau4R;AP;TDP-43cKO* mice but had begun to spread to the olfactory bulb (OB), thalamus (TH), hypo-thalamus (HTH), midbrain (MB) as well as the brain stem (BSM) (**Fig. 2a, d, second column; Sup Fig. 4a, second column & 5b**). As compared to *Tau4R;AP* littermates, 24-months old *Tau4R;AP;TDP-43cKO* mice harbored marked increase of tau burden in these brain regions (**Fig. 2b, e, second column; Sup Figs. 4b, 5b & 6a**). These findings establish that early TDP-43 dysfunction accelerates the tau pathology in *Tau4R;AP;TDP-43cKO* mice. To determine whether in *Tau4R;AP;TDP-43cKO* mice the emergence and spread of TDP-43 cytoplasmic aggregates is dependent on the spread of tauopathy, we monitored the pattern of tau aggregation as it spreads to different brain regions during aging. Remarkably, the emergence and spread of TDP-43 pathology depends on the spread of the accelerated tauopathy observed in *Tau4R;AP;TDP-43cKO* (**Fig. 2c, f, second column; Sup Fig. 2 a-d, second column; Sup Fig. 4a-b, second column**). These results establish that the emergence and spread of pTDP-43 proteinopathy is dependent on the spread of tau pathology.

**Figure 5:**
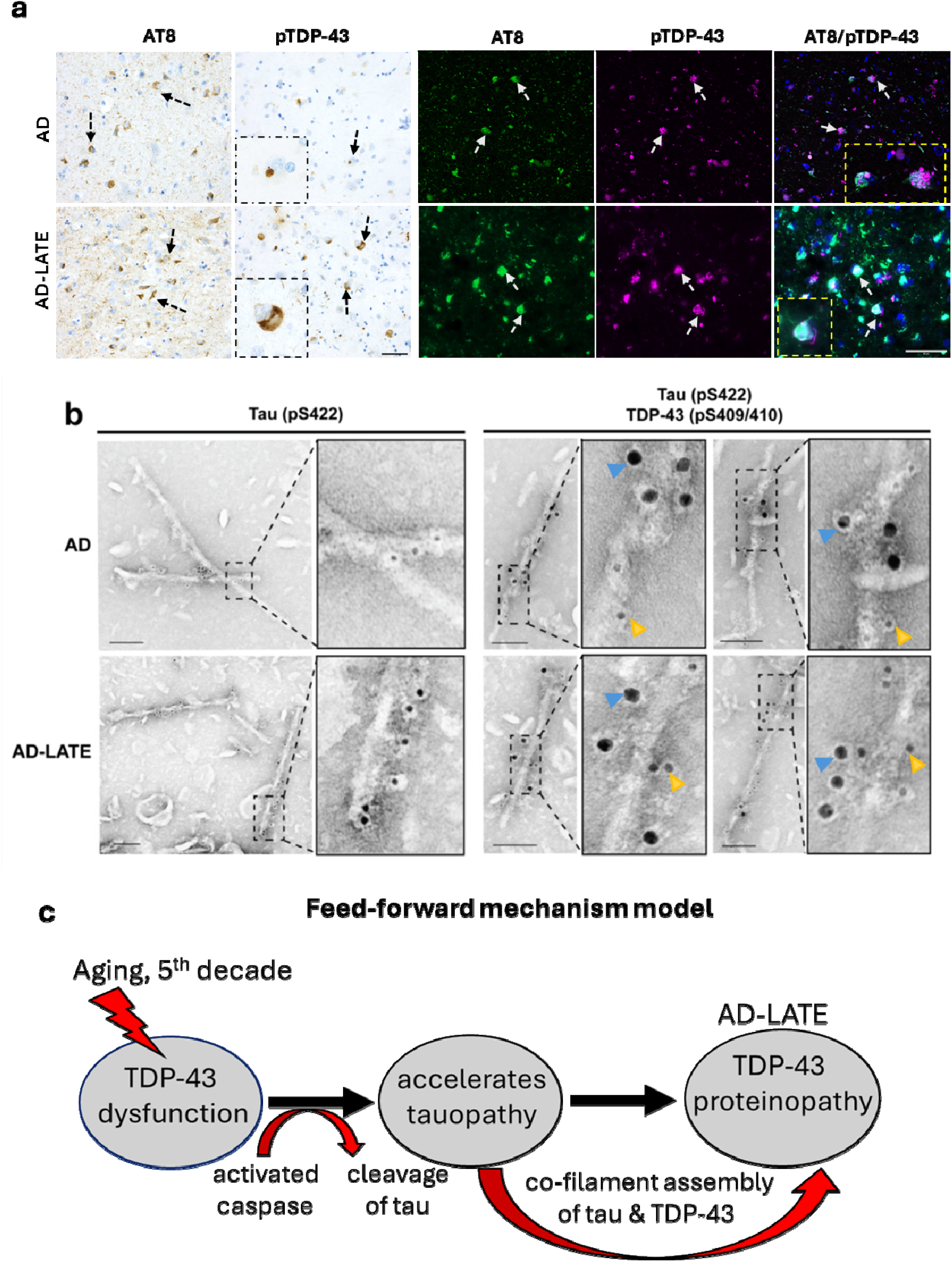
Co-filament assembly of tau and TDP-43 occurs in brains of AD and AD-LATE. (**a**) Immunohistochemical and neuropathological analysis using antibody AT8 (pSer202, pThr205) in of amygdala from AD and AD-LATE brains, Scale bar, 20 μm. Double immunofluorescence shows the co-localization of pTau (AT8) and pTDP-43 (Arrow heads show co-localization of pTau (AT8) and pTDP-43 (cytoplasmic small inclusions) in AD and with mature cytoplasmic pTDP-43 aggregation in AD-LATE brains (Arrow heads show immunoreactivity of AT8 and pTDP-43, Scale bar, 50 μm). (**b**) Double labeling immuno-EM analysis of sarkosyl-insoluble brain extracts derived from AD and AD-LATE amygdala samples using primary antibodies against tau (pS422) and TDP-43 (pS409/410) and goat anti-rabbit 6nm (yellow arrowheads) gold-conjugated and goat anti-mouse 12nm (blue arrowheads) gold-conjugated secondary antibodies (scale bar, 100nm). (**c**) Schematic diagram showing proposed model of feed-forward mechanism of how TDP-43 dysfunction accelerates tauopathy may cause emergence of TDP-43 pathology and its dysfunction exacerbate neuron loss in time dependent manner AD-LATE

### TDP-43 dysfunction accelerates tauopathy and TDP-43 proteinopathy to exacerbate neuron loss

That TDP-43 dysfunction occurs during aging or pre-symptomatically in human disease (*5–7*) coupled with our recent finding that loss of TDP-43 can facilitate and accelerate the pathological conversion of endogenous tau (*29*) suggest that early TDP-43 dysfunction accelerates tauopathy dependent TDP-43 proteinopathy thereby exacerbates neuron loss and brain atrophy in AD-LATE (*18–19*).

As expected, depletion of TDP-43 led to selective neuron loss of hippocampal Cornus Ammonis (CA) 2/3 region (**Fig. 3a, upper panel second column**). As compared to wild-type mice, no neuronal loss was observed in the hippocampus of *Tau4R;AP* by age of 20 months but TDP-43 depletion exacerbated brain atrophy and neuron loss not only in CA2/3 but also in dentate gyrus (DG) and CA1 subfields (**Fig. 3a, upper panel; Sup Figs. 7 & 8a**). Furthermore, aging accelerated brain atrophy and neuron loss in *Tau4R;AP* and *Tau4R;AP;TDP-43cKO* mice (**Fig. 3a, lower panel**). To determine neuron loss, we quantified the number of NeuN positive neurons in CA1, CA2/3, and DG. As expected, TDP-43 deletion led to CA2/3 neuron loss (**Fig. 3b**); in contrast, by 20 months neurons in DG and CA1 of *Tau4R;AP;TDP-43cKO* mice are also markedly decreased (**Fig. 3c-d**). As compared to neuron loss restricted to CA1 region in *Tau4R;AP* mice, a marked exacerbation of neuron loss is observed in CA1, CA2/3 and DG in *Tau4R;AP;TDP-43cKO* mice by 24 months (**Fig. 3e**). These results are consistent with a model for the selective vulnerability of neurons to TDP-43 dysfunction in the mouse hippocampus: CA2/3 neurons are most vulnerable, whereas CA1 or DG granular neurons, respectively, are vulnerable to the co-pathology of tau and TDP-43 or co-pathology of amyloid-β and tau (**Sup Fig. 7a-d & 8a-e**).

To establish whether early TDP-43 dysfunction accelerates the tau pathology in *Tau4R;AP;TDP-43cKO* mice, we assessed the tau burden using an antibody recognizing misfolded tau. Remarkedly, as compared to *Tau4R;AP* mice, we observed that early TDP-43 dysfunction facilitated greater burden of tau pathology in *Tau4R;AP;TDP-43cKO* mice (**Fig. 3f-g; Sup Fig. 4-8**). The aggregation form of tau was confirmed by extracting sarkosyl-insoluble tau aggregates (P3 fraction) that are detectable by immunoblotting using anti-pTau antibody (AT8) (**Fig. 3h**). These observations establish that TDP-43 dysfunction accelerates the pathological conversion of endogenous tau to drive TDP-43 proteinopathy and exacerbated neuron loss in *Tau4R;AP;TDP-43cKO* mice, providing the opportunity to determine the pathogenic mechanism of tauopathy-dependent TDP-43 proteinopathy.

As we demonstrated, depletion of TDP-43 accelerates tauopathy conversion through amplification of caspase 3 activation-mediated tau cleavage in Tau4R mice (*29*). Therefore, we confirmed that depletion of TDP-43 amplifies the activation of caspase 3-mediated cleavage of tau levels in the *Tau4R;AP;TDP-43cKO* mice (**Fig. 3i; Sup Fig. 6b & Sup Fig 7e, f**), suggesting that TDP-43 dysfunction accelerates the pathological conversion of tau through amplification of caspase 3-mediated cleavage of tau.

### Tauopathy primes dysfunction and co-filament assembly of TDP-43

To determine how tauopathy promotes TDP-43 cytoplasmic aggregation, we asked whether tau filaments can facilitate the aggregation of TDP-43. Biochemical and immunohistochemical evidence from human AD brains (*38, 41*) and in our mouse model (**Sup Fig. 3c**) suggest that pTDP-43 and phosphorylated tau co-aggregate, an event that could be preceded by the co-filament assembly of tau and TDP-43. To assess the presence of both TDP-43 and tau filaments, we performed double labeling immunoelectron microscopy (immuno-EM) on sarkosyl-insoluble filament extracts derived from 20- and 24-month-old *Tau4R-AP* and *Tau4R-AP;TDP43cKO* brains using primary antibodies recognizing pTau (S422) or pTDP-43 (S409/410) and secondaries conjugated to 6nm and 12nm gold particles, respectively. At 20 months of age, *Tau4R-AP;TDP-43cKO* but not *Tau4R-AP* mice exhibit mostly singly labeled pTau filaments (**Fig. 4b, inset view in right column**) but individual pTDP-43 immunoreactive filaments are essentially non-existent or rarely observed (**Sup Fig. 9a**). Unexpectedly at this earlier time point, in *Tau4R-AP;TDP-43cKO*, but not *Tau4R-AP,* mice, filaments co-labeled with both pTau and pTDP-43 antibodies were evident (**Fig. 4e, inset view in right column**). Notably, preceding co-filament assembly, we observed cryptic exon (*Unc13a & Ift81*) inclusion (**Fig. 4a, first column**) in brains of *Tau4R;AP* mice. By 24 months of age, in brains of *Tau4R-AP* mice, we observed the emergence of co-filament assembly of tau and TDP-43 associated with cryptic exon inclusions (**Fig. 4c-d**). For both *Tau4R-AP and Tau4R-AP;TDP-43cKO* mice, most filaments were pTau immunoreactive (**Fig. 4d, f, inset view in right column**) accompanied by a few pTDP-43 labeled filaments (**Sup Fig. 9a**). Furthermore, we also observed that these co-labeled filaments of tau-TDP-43 are obviously present in the brain of *Tau4R-AP;TDP-43 cKO* at 24 months of age (**Fig. 4f, inset view in right column**). No labeled filaments were observed in control genotype mice (**Sup Fig. 10**).

These novel observations of ptau and pTDP-43 co-labeled filaments in our mouse models led us to probe for the existence of such filaments in brains of AD and AD-LATE cases. Co-localization of tau and TDP-43 aggregates has been demonstrated in AD-LATE brains (*38, 42–44*). We first verified this observation by double immunofluorescent staining, which revealed that clear aggregates of phosphorylated tau and TDP-43 co-localized in the AD-LATE brains (**Fig. 5a, lower panel**), but interestingly we also observed small pTDP-43 inclusions co-exists with tau aggregates in AD brains (**Fig. 5a, upper panel**). This observation suggests that in AD brains, tau pathology may have induced small pTDP-43 inclusions but did not evolve to mature aggregates. Furthermore, to investigate the presence of co-filaments comprised of tau and TDP-43, we extracted filaments from four AD and AD-LATE amygdala samples and performed double labeling immuno-EM analysis. Like what we observed in our mouse models, most filaments were phosphorylated tau immunoreactive (**Fig. 5b**, **first column with inset view in right**). In all cases of AD-LATE, several co-labeled filaments were identified (**Fig. 5b**, **lower panel, right two columns with inset view, Sup Fig. 11**). Surprisingly, we also observed the co-filament assembly of tau and TDP-43 across all AD cases (**Fig. 5b, upper panel, right two columns with inset view, Sup Fig. 12**). As controls, we demonstrated that no labeled filaments were present in control and Parkinson’s Disease (PD) cases (**Sup Fig. 12d-e**). To verify the specificity of the antibodies, we included a secondary only control to show that no filaments were labeled (**Sup Fig. 9b**).

Our data and previous human studies together support our feed-forward model of mechanism (**Fig. 5c**); where early TDP-43 dysfunction can accelerate tauopathy in human aging populations (*6*) may be through activated caspase-mediated tau cleavage as observed in mouse model (**Fig. 2-3; Sup Figs. 6b & 7e-f**). These findings strongly support the view that TDP-43 dysfunction accelerates tauopathy through activated caspase-cleaved tau which in turn primes TDP-43 proteinopathy through co-filament assembly of tau and TDP-43 leading to worsened neurodegeneration occurring in AD-LATE.

## Discussion

A central problem in resolving the complexity of AD pathogenesis is how the co-pathology of TDP-43, found in a majority of ADRD (*18–19, 23–26*), contributes to the exacerbated neuron loss. We addressed this question by using our previous established tau seeding mouse model (*Tau4R;AP* mice) exhibiting the canonical pathologies of Aβ and tau and demonstrated that tauopathy is necessary to drive TDP-43 proteinopathy to exacerbate loss of vulnerable neurons. Given our previous that TDP-43 dysfunction is an aging risk factor to increase the burden of tau pathology (*6*), our current findings are consistent with the idea that TDP-43 dysfunction accelerates tauopathy to drive exacerbated cell loss. These novel findings would thus support a feed-forward model whereby TDP-43 dysfunction accelerates tauopathy and in turn tauopathy drives TDP-43 cytoplasmic aggregation and dysfunction as judged by inclusion of TDP-43 cryptic exons. This feed-forward model is consistent with the observations in our mouse model (**Figs. 2-3**) as well as studies in longitudinal aging and diseased brains showing early nuclear clearance of TDP-43 can accelerate the burden of tau pathology (*6*), which could prime TDP-43 proteinopathy (**Figs. 1-3**) in AD-LATE (*6, 20, 38*). By correlating this novel finding of co-filament assembly of tau and TDP-43 that precedes the emergence of TDP-43 cytoplasmic aggregation, we provide a potential pathogenic mechanism whereby tauopathy facilitates the maturation of cytoplasmic TDP-43 aggregates in AD-LATE (**Fig. 5c**). The disease relevance of such a view is supported by our finding of co-filament assembly of tau and TDP-43 in brains of cases of AD or AD-LATE. We thus hypothesize that TDP-43 dysfunction is a major risk factor occurring in AD-LATE individuals that would accelerate tauopathy, providing sufficient time to permit the emergence of TDP-43 proteinopathy. Future spatial transcriptomic approaches using mouse models and human brain tissues will be necessary to further clarify the molecular mechanisms of tauopathy-dependent TDP-43 proteinopathy.

A recent *in vitro* study demonstrating the direct interaction of tau and TDP-43 co-filament assembly (*41*), would be consistent with the idea that tauopathy could prime TDP-43 proteinopathy in AD-LATE. Since heterotypic filament comprised of A11 and TDP-43 is observed in FTLD-TDP type C (*45*), future cryo-EM analysis will be necessary to resolve whether the co-filament assembly of tau and TDP-43 we observed in AD and AD-LATE and their mouse models are homo- or heterotypic filaments. In view of our finding of co-filament assembly of tau and TDP-43, recent studies showing chronic traumatic encephalopathy (CTE) like tau fold in AD-LATE brains (*46*) could be reconciled by tau and TDP-43 co-filament assembly. Solving the structure of tau and TDP-43 co-filament will be crucial for identification of novel therapeutic targets for preventing the emergence of TDP-43 cytoplasmic aggregates to attenuate neuron loss in AD-LATE patients. With the advent of alphafold-3 (*47*), prediction algorithms may identify small molecules localized selectively to the interface formed by tau and TDP-43 to disrupt their co-filament assembly.

Moreover, our finding in *Tau4R;AP;TDP-43cKO* mouse model that early loss of TDP-43 accelerates tauopathy led to greater neuron loss (**Fig 3**), suggests that therapies designed to restore TDP-43 function would attenuate neurodegeneration for AD-LATE individuals. Validating our gene therapy strategy to restore TDP-43 function (*2, 48–50*) as a monotherapy or in combination with a tau drug, such as an ASO against tau (currently in Phase II clinical trials), in our *Tau4R;AP* mice should be instructive.

## Acknowledgment

We thank T. Melnikova, B. Kim, C. Kim, and Y. Zhao for technical support. This work was supported in part by NIH grants (R01NS095969, R33/R61NS115161 and UG3/UH3R61NS115608161) to PCW.

## Authors contribution

M.S.B. conceived, designed and coordinated the studies, performed the histological, biochemical, statistical analysis, and wrote the manuscript. G.D.B. performed Sarkosyl-insoluble extraction, immuno-EM studies, and wrote the manuscript. A.P.M. performed BaseScope analysis of cryptic exons, co-detection with protein, and helped in quantifications. X.K.C. has helped in immunohistochemistry and neuronal quantifications, and F.L helped in Sarkosyl-insoluble extractions. S.R. performed brain sectioning and technical help. P.C.W. conceived and designed studies and edited the manuscript. T.L. has edited the manuscript. J.C.T. neuropathologist BRC, at Hopkins provided human frozen brain materials and sections, and all authors read and approved the final manuscript.

## Methods

### Mouse Models

All the animal experiments were conducted as per the regulations of the Animal Care and Use Committee at Johns Hopkins University School of Medicine in accordance with the laws of the State of Maryland and the United States of America. We used the conditional knockout mouse model of TDP-43 by generating CaMKIIα-CreER;Tardbp^f/f^ (TDP-43 KO) mice with loxP sites flanking Tardbp exon 3 (*30*), creating a tamoxifen-induced recombination in excitatory forebrain neurons in adult mice. Through a three-stage breeding strategy (**Supp Fig. 1a**), CaMKIIα-CreER;Tardbp^f/f^ mice bred with tTA;Tau4R;APP (Tau4R;AP) line generated previously (*27*) and a cohort of tTA;Tau4R;CaMKIIα-CreER;Tardbp^f/f^ (Tau4R;AP;TDP-43 cKO) mice on a C57BL/6J background, as well as littermate controls. Animals were genotyped at weaning, and mice were subsequently housed with one littermate of each genotype (5 mice/cage) when possible. Oral administration of tamoxifen citrate through diet was done in all animals (Harlan Teklad) at an average amount of 40 mg/kg/day for a 4-week period, beginning at 12 months-of-age. Studies in transgenic Tau4R;AP;TDP-43 cKO mice and littermate controls were carried out in brain tissue dissected from 20- or 24-month-old mice.

The TDP-43 cKO;AP mice cohort was administered tamoxifen to delete TDP-43 at the age of 6-months and analysed at 15 and 22-month (**Sup Fig. 3**)

For the AAV-PhP.eB-hTau4R/GFP injection model, we injected wild-type mice with AAV-PhP.eB-hTau4R/GFP stereotaxic surgery method.

### Human Samples

The postmortem brain tissues used in the present study were provided by the Johns Hopkins Brain Resource Center. For histological analysis, formalin-fixed paraffin-embedded (FFPE) tissue Sections. (10 μm) of the Amygdala were obtained from 3 pathologically confirmed AD and AD-LATE cases with Braak neurofibrillary stages II-VI. Details of human cases used for histology and Sarkosyl-insoluble filament study by immune-electron microscopy are available in respective tables (**Table S1 and S2**). The cohort (n = 6) included 3 males and 3 females ages 61 to >90 years. All AD and AD-LATE subjects had been prospectively recruited, clinically characterized by the Johns Hopkins Alzheimer’s Disease Research Center (ADRC), and underwent neuropathologic postmortem examination excluding Lewy body disease or non-AD tauopathies. The clinical and autopsy components of this study were approved by the Johns Hopkins Medicine IRB.

### Histology and immunohistochemistry

Transgenic Tau4R;AP and Tau4R;AP;TDP cKO mice and their age matched littermates were sacrificed at 20 or 24 months of age for analysis. Injected mice were sacrificed 10 months post AAV-hTauRD injection. Mice were anesthetized with isoflurane and transcardially perfused with ice-cold PBS. Brains were dissected, postfixed in 4% PFA for 24 hours, embedded in paraffin and then sectioned with a microtome into 10µm thick sagittal sections.

For immunohistochemistry, the slides were deparaffinized and antigen retrieval was performed using 10mM sodium citrate buffer to efficiently expose the epitope to the antibodies. Nonspecific binding of antibodies was eliminated by incubating with blocking buffer (1.5% normal goat serum in PBS with 0.1% Triton-X) for 1 hour. Slides were then incubated in 0.3% H2O2 for 30 minutes to quench endogenous peroxidase. Primary antibodies were prepared in blocking buffer and applied overnight at RT followed by incubation with biotinylated secondary antibody for a 1 hour. Peroxidase labeled ABC reagent (Vector Laboratories) was applied for 30 minutes followed by signal development using 3,3 diaminobenzidine (DAB) (Vector Laboratories). Slides were counterstained with hematoxylin, dehydrated, cleared and mounted.

The following primary antibodies were used: antiserum against NeuN (1:1,000; MAB377, Merck), TDP-43 N-terminus (1:1,000; 10782-2-AP, ProteinTech), phosphorylated tau AT8 (1:1,000; MN1020, Invitrogen), TDP-43 S409 (1:1000; CAC-TIP-PTD-P03; Cosmo Bio), TDP-43 S409/410 (1:1000; CAC-TIP-PTD-M01; Cosmo Bio), cleaved Caspase 3 (1:2000; D3E9; Cell Signaling Technology), cleaved tau (TauC3) (1:200; AHB0061, Invitrogen).

The numbers of healthy neurons in the CA1, CA2/3, and DG regions were counted manually using ImageJ. The minimum three sections from three different planes were used for quantification.

A minimum of 2-3 sections for each mouse were used to quantify DAB staining, images were color deconvoluted, and detection threshold was set manually by comparison to the original signal. The measured relative DAB intensity values are plotted in the corresponding graph. For TauC3 quantification, we manually counted the TauC3 (considering signal in cell body) positive cells in hippocampal slices. We used 2-3 slices from each mouse to quantify the TauC3 positive cells in different groups.

AT8 and pTDP positive cells were quantified manually using Image J.

### Immunofluorescence Staining

For immunofluorescent staining of mouse and human brain tissues, to perform immunofluorescence staining on tissue, the paraffin brain sections were deparaffinized and antigen retrieval was performed by boiling for 4 minutes in 10mM sodium citrate buffer to expose the epitope to the antibodies. Nonspecific binding of antibodies was eliminated by incubating with blocking buffer (1.5% normal goat serum in PBS with 0.1% Triton-X) for 1 hour. After blocking, the primary antibodies against phosphorylated tau AT8 (1:1,000; MN1020, Invitrogen) and pTDP-43 S409 (1:1000; CAC-TIP-PTD-P03; Cosmo Bio) (1:1000; 79327 were incubated overnight in humid chambers. Unbound antibodies were washed out and incubated with respective secondary conjugated fluorophores (Alexa Fluor 488 & Alexa Fluor 594) and mounted with antifade DAPI. The Zeiss Apotome Inverted Fluorescence Microscope (Zeiss, Germany) was used for imaging.

### Intraparenchymal stereotaxic injection

Mice were deeply anesthetized with isoflurane and fixed in a stereotaxic frame. The scalp skin of the animal was swabbed with povidone iodine and the skull was exposed via an incision. The skull was cleaned with hydrogen peroxide to allow for a craniotomy to be made above the right hippocampus using coordinates for dorsal hippocampus (2 mm frontal to lambda, 2 mm lateral to midline, and 1.5 mm depth) from the skull surface at the rate of 1µL/min as standardized before (*31*). Unilateral injections were performed using a syringe (Hamilton) containing 5 µL of 2x1013 vg/mL AAV-PhP.eB-GFP or hTauRD. Upon completion, the needle was slowly withdrawn for 3 minutes. The skin was stapled and covered with 2% chlorohexidine to heal the incision.

### Sarkosyl extraction

Brain tissue was weighed and put in Beckman polycarbonate thick-wall centrifuge tubes. 10X volume of homogenization buffer (TBS pH: 8.0, 50mM Tris base, 274mM NaCl, 5mM KCl, 1X protease inhibitor, 1X phosphatase inhibitor and 1mM PMSF) was used to homogenize brain using 15 strokes in a 1-3ml glass dounce. Homogenate was centrifuged at 28,000 rpm for 20 min at 4°C. Supernatant was aliquoted to new eppendorf tubes as S1 (TBS-SOLUBLE) and stored at –80oC. Pellet (P1) was homogenized with extraction buffer (10mM Tris, pH 7.4, 0.8M NaCl, 10% sucrose, 1mM EGTA, 1mM PMSF), ∼5x volume of wet weight of the original tissue. Homogenate was centrifuged at 22,000 rpm for 20 minutes at 4°C. The pellet (P2) was stored at –80°C and the supernatant (S2) was transferred to a new Beckman polycarbonate thick-walled tube. The supernatant was incubated with 1% Sarkosyl and rotated for 5 min at room temp, then shaken at 37°C for 1 hour. Samples were then centrifuged at 55,000 rpm for 1 hour at 4°C. Supernatant was aliquoted into new eppendorf tubes as S3 (SARKOSYL-SOLUBLE) and kept at –80°C. Pellet was resuspended in TE buffer (10mM Tris, pH 8.0 and 1mM EDTA) by pipetting up and down thoroughly and labeled as P3 (SARKOSYL-INSOLUBLE) then stored at –80°C.

### Immunoblotting

For protein blot analysis of Sarkosyl insoluble fraction, samples (S3,P3) were resolved on 4–12% Bis-Tris SDS–PAGE gels and then transferred to polyvinylidene difluoride membranes (Invitrogen, Carlsbad, CA). After blocking with 5% skimmed milk, the membranes were probed with the following antibodies: phosphorylated tau AT8 (1:1,000; MN1020, Invitrogen), pTDP-43 S409/410 (1:1000; CAC-TIP-PTD-M01; Cosmo Bio) for overnight incubation. Next day, primary unbound antibody washed, then membrane was incubated with secondary antibodies HRP-conjugated for 1 hr. Immunoblots were developed using enhanced chemiluminescence method (Millipore Corp., MA).

### Immuno-Electron microscopy

Samples were absorbed to 400 mesh carbon coated nickel grids (presoaked in dH2O and blot dried) for 2 min (with blot). Grids were transferred (anti-capillary forceps) to 1% NGS 20 mM TBS 150 mM NaCl for 20 min, then incubated in primary antibody diluted 1:40 in blocking solution for 3 hours under a humidity chamber. Controls were incubated in blocking only. Grids were then transferred to blocking solution for 10 min, quickly washed in three drops of TBS and floated on secondary antibody solution (goat anti-mouse 12nm gold or goat anti-rabbit 6nm gold, 1:40 dilution) for 2 hours under a humidity chamber. Following a 10 min TBS drop and 3 quick TBS passes, grids were fixed in 1% glutaraldehyde in 100 mM cacodylate for 5 min, rinsed for 1 min in dH2O, then negatively stained with 2 drops of fresh and filtered uranyl acetate (aqueous). Grids were blot dried and viewed 24 hours later on a Hitachi H-7600 operating at 80 kV. Images were captured with an AMT XR80 CCD (8 megapixels).

### BaseScope-ISH and Co-detection assay

RNA in situ hybridization was carried out using the BaseScope Detection Reagent v2-RED Assay Kit (Advanced Cell Diagnostics, Inc., Ref#323900) according to the manufacturer’s protocol. Transcripts containing cryptic exon splice sites in *Unc13a* and *Ift81* were detected with custom 3zz BaseScope probes (BA-Mm-Unc13a-O1-2EJ-C1, Ref# CAT#1182491-C1 and BA-Mm-*Ift81*-E17-intron17-NJ, Ref#712201 respectively). RNA integrity and assay performance were evaluated using positive and negative control probes (BA-Mm-Ppib, Ref#701071 and BA-DapB, Ref#701011). Briefly, consecutive paraffin-embedded tissue sections (10 μm thick) were deparaffinized and sequentially pretreated with hydrogen peroxide, target retrieval buffer, and Protease IV. The sections were then hybridized with target probes in a HybEZ II oven (Advanced Cell Diagnostics, Inc.) at 40°C for 2 hours. Following signal amplification, slides were counterstained with hematoxylin. Images were captured using a Zeiss Apotome inverted brightfield microscope (Zeiss, Germany).

Co-detection assay was performed on serial sections to enable the simultaneous detection of transcripts containing cryptic exons (*Ift81*) and TDP-43 proteins within the same cells using RNA-protein Co-detection Ancillary Kit (Advanced Cell Diagnostic Inc. Ref#323180) according to the manufacturer’s instructions. Briefly each section was subjected the pretreatment, followed by overnight incubation with the primary antibody at 4°C. The primary antibody was then washed off the following day, and the sections were post-fixed in 10% neutral buffered saline for 30 min at RT. Sections were then washed and proceeded with the BaseScope assay as described. After the detection step, sections were treated with co-detection blocker and incubated with the secondary antibody, followed by the detection of protein signal using 3,3’-diamino benzidine (DAB) staining. The sections were counterstained in 50% Gill’s hematoxylin solution No. 1 (Electron Microscopy Sciences, Ref#26030) and mounted using EcoMount (Biocare Medical, Ref# EM897). Images were captured using a Zeiss Apotome Inverted Brightfield Microscope (Zeiss,Germany).

### Statistical analysis

Graphs were generated using GraphPad Prism. Individual data points are also shown. One-two way analysis of variance (one-way ANOVA) for multiple comparisons was performed using the GraphPad Prism software (La Jolla, CA, USA). In all tests, values of p<0.05 were considered significant. The number of biological replicates and details of statistical analyses are provided in the figure legends.

---

**Table S1:**
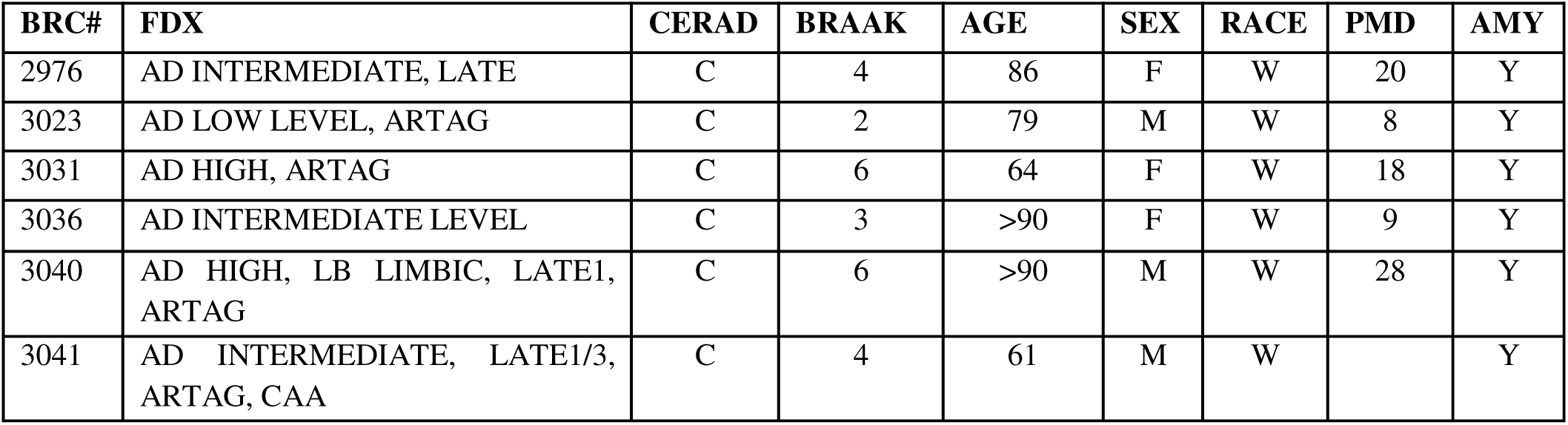
Human brain tissues used for immunohistochemistry.

**Table S2:**
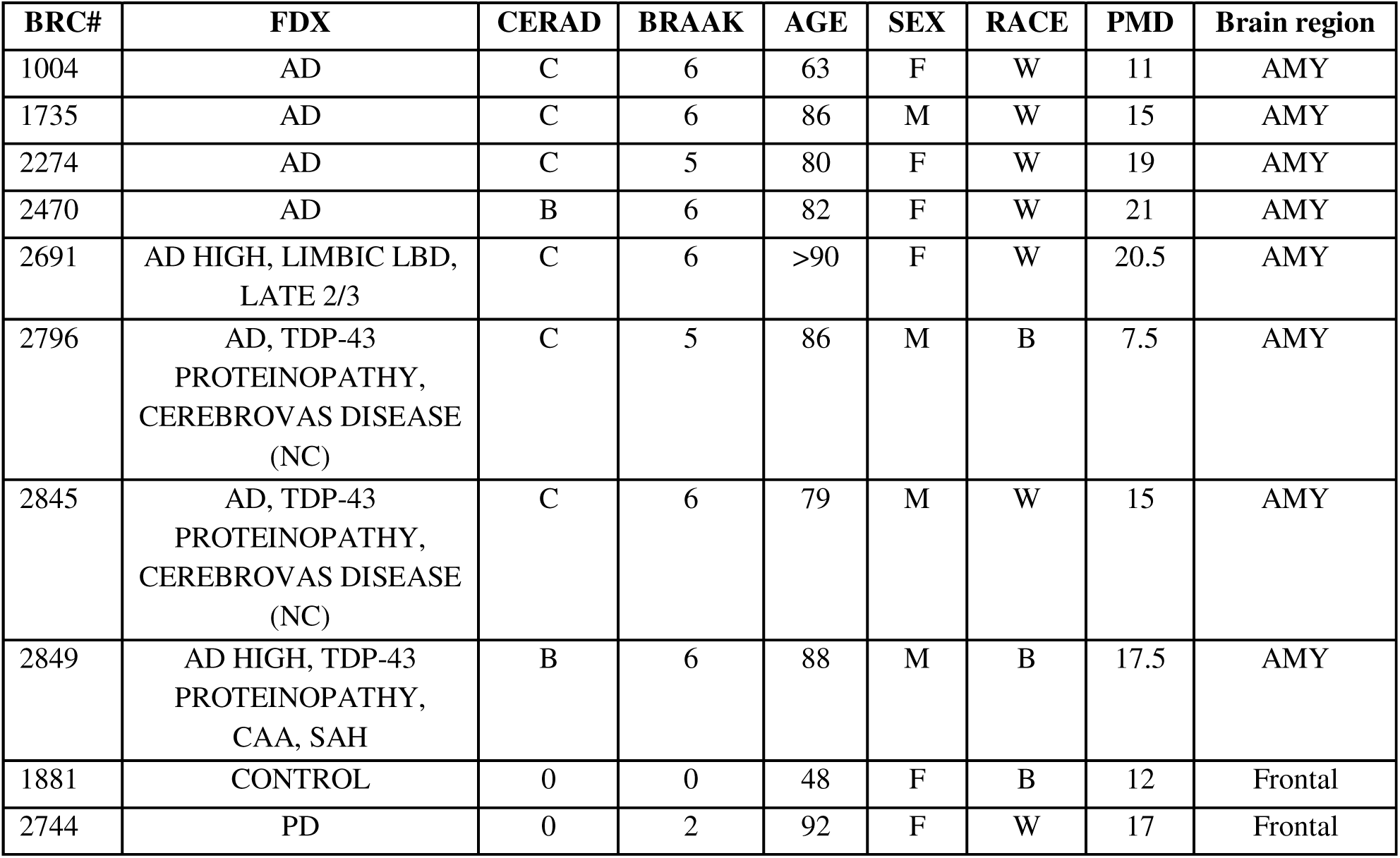
Human brain tissues used for immuno-electron microscopy.

## Supplementary Figures

**Sup Fig. 1:**
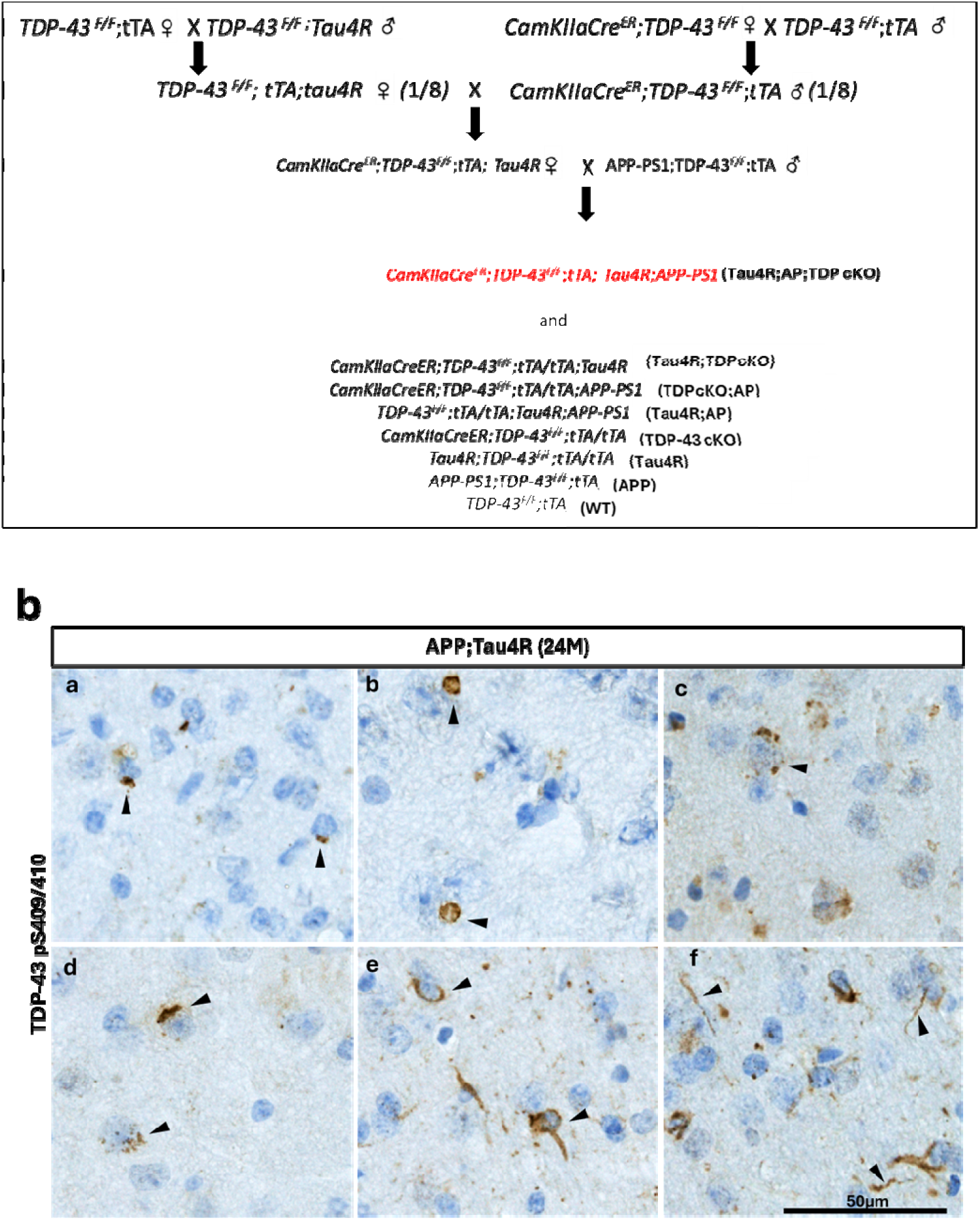
**(a)** Crossbreeding strategy to generate mouse model Tau4R;AP; TDP-43 cKO and respective control genotypes. **(b)** Immunohistochemical analysis of brains of 24-month-old Tau4R;AP mice using antibody specific to pTDP-43 409/410 to detect different cytoplasmic aggregation patterns in brain (Arrow heads show the pTDP-43 cytoplasmic aggregations, scale bar 50um).

**Sup Fig. 2:**
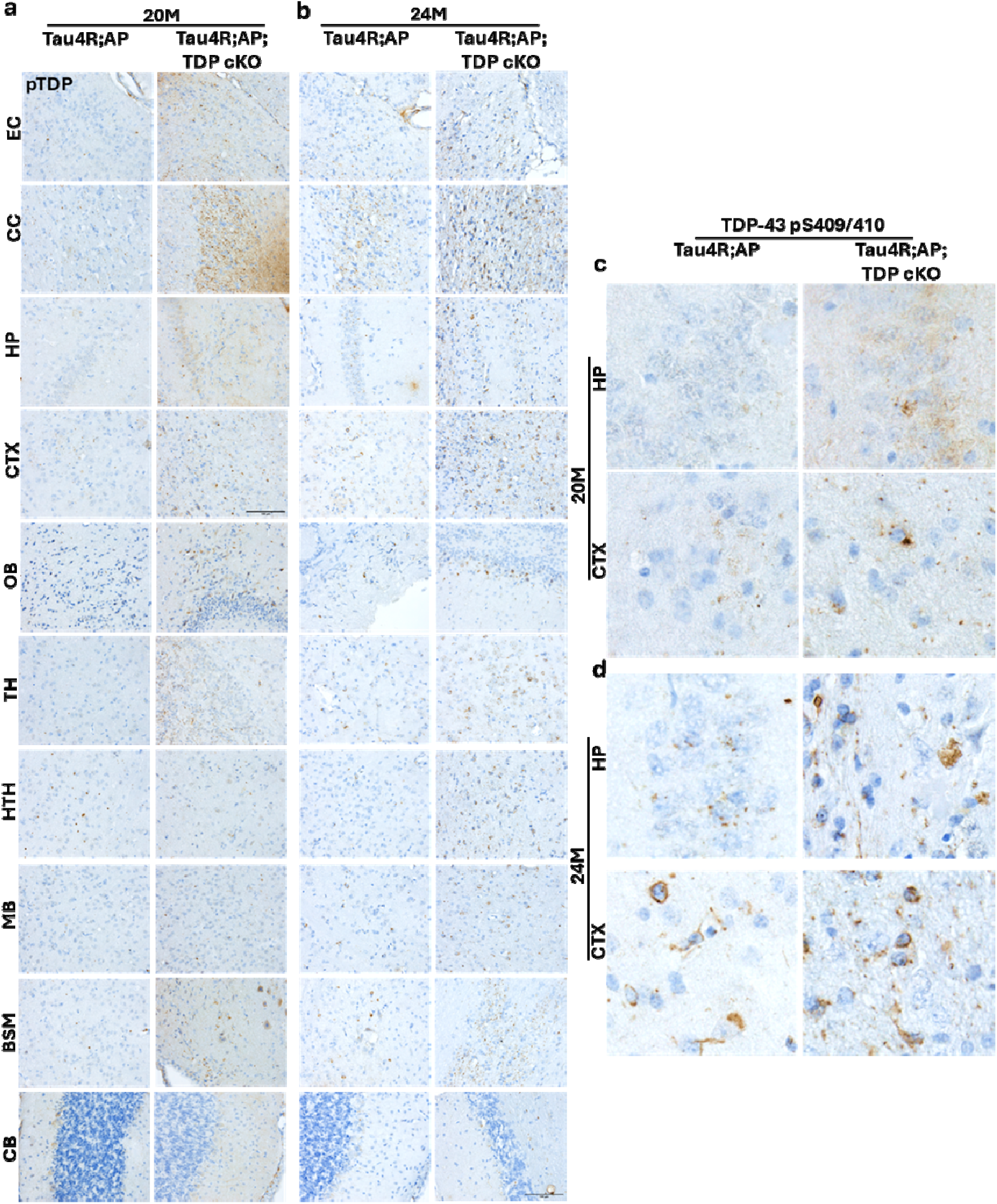
Loss of TDP-43 function accelerates the pTDP-43 pathology in the brain of *Tau4R;AP;TDP cKO* mice in age dependent manner: (**a-b**) Immunohistochemical analysis extended panel of brains of 20 and 24-month-old Tau4R;AP and Tau4R;AP;TDP cKO using specific antibody TDP-43 pS409/410 to show the TDP-43 pathology emergence in tau spread directed manner in brain regions. **(c-d)** Immunohistochemistry of pTDP-43 panel shows enlarged views of pTDP-43 inclusions in HP and CTX of 20- and 24-months old respectively of Tau4R;AP and Tau4R;AP;TDP cKO mice.

**Sup Fig 3:**
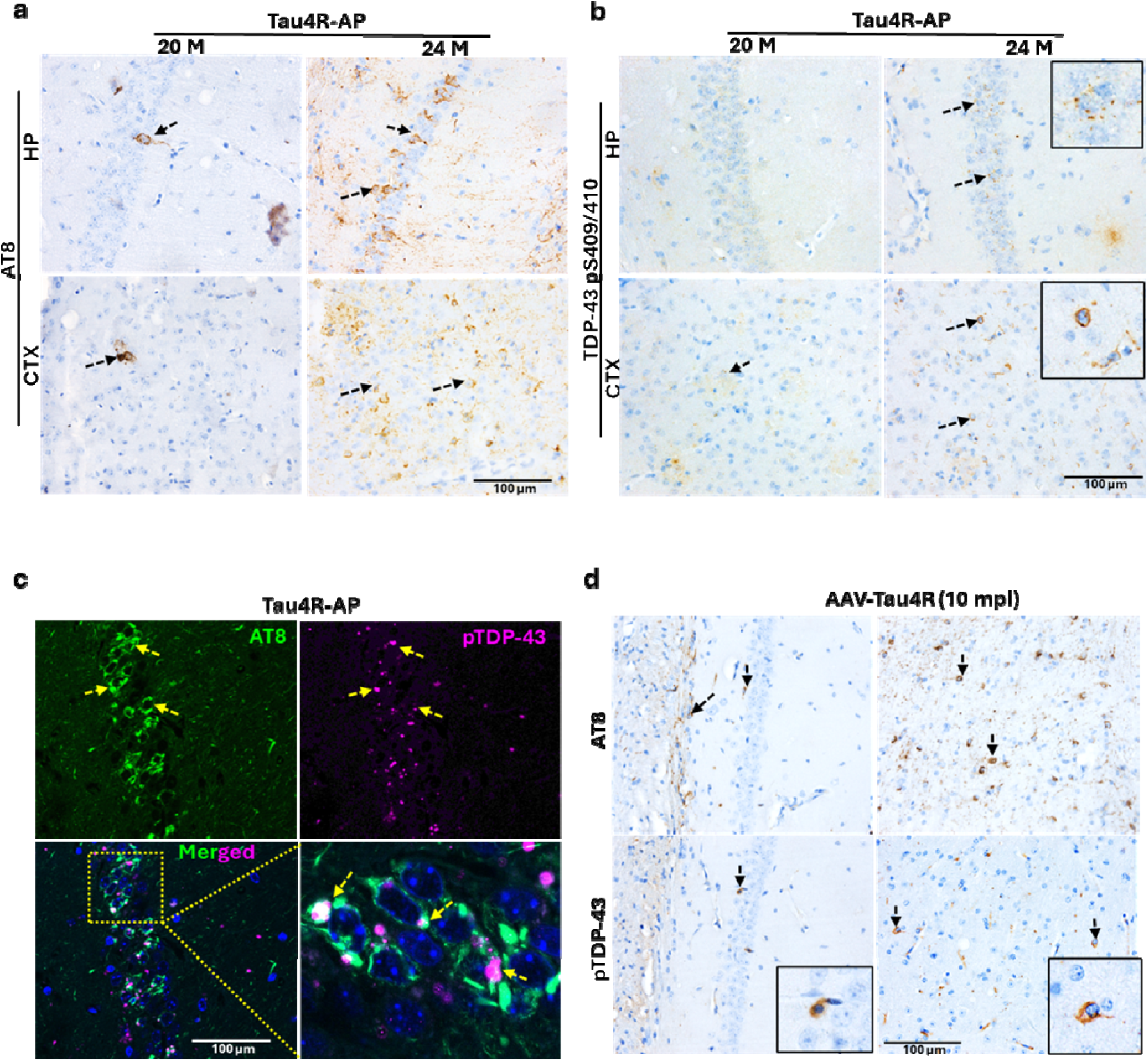
Pathological conversion of tau is sufficient to induce TDP-43 proteinopathy. (**a**) Immunohistochemical (IHC) analysis of the hippocampus and cortex using antibody specifi to endogenous phosphorylated tau AT8 (pSer202, pThr205) in 20- and 24-month-old Tau4R;AP mice (n=3), Scale bar, 100 μm. (**b**) Immunohistochemical analysis of the hippocampus and cortex using antibody specific to pTDP-43 in 20- and 24-month-old Tau4R;AP mice, (n=3), Scale bar, 100 μm. (**c**) Double immunofluorescence analysis shows the co-localization of pTau (AT8) and pTDP-43 (Arrow heads show co-localization of AT8 and pTDP-43) in *Tau4R;* A*P* mice (Scale bar, 100 μm). (**d**) IHC analysis showing the pathological conversion of endogenous tau using AT8 antibody in hippocampus and thalamus (upper panel), derived pTDP-43 aggregation (lower panel) (Arrow heads show immunoreactivity of AT8 and pTDP-43, Scale bar, 100 μm).

**Sup Fig. 4:**
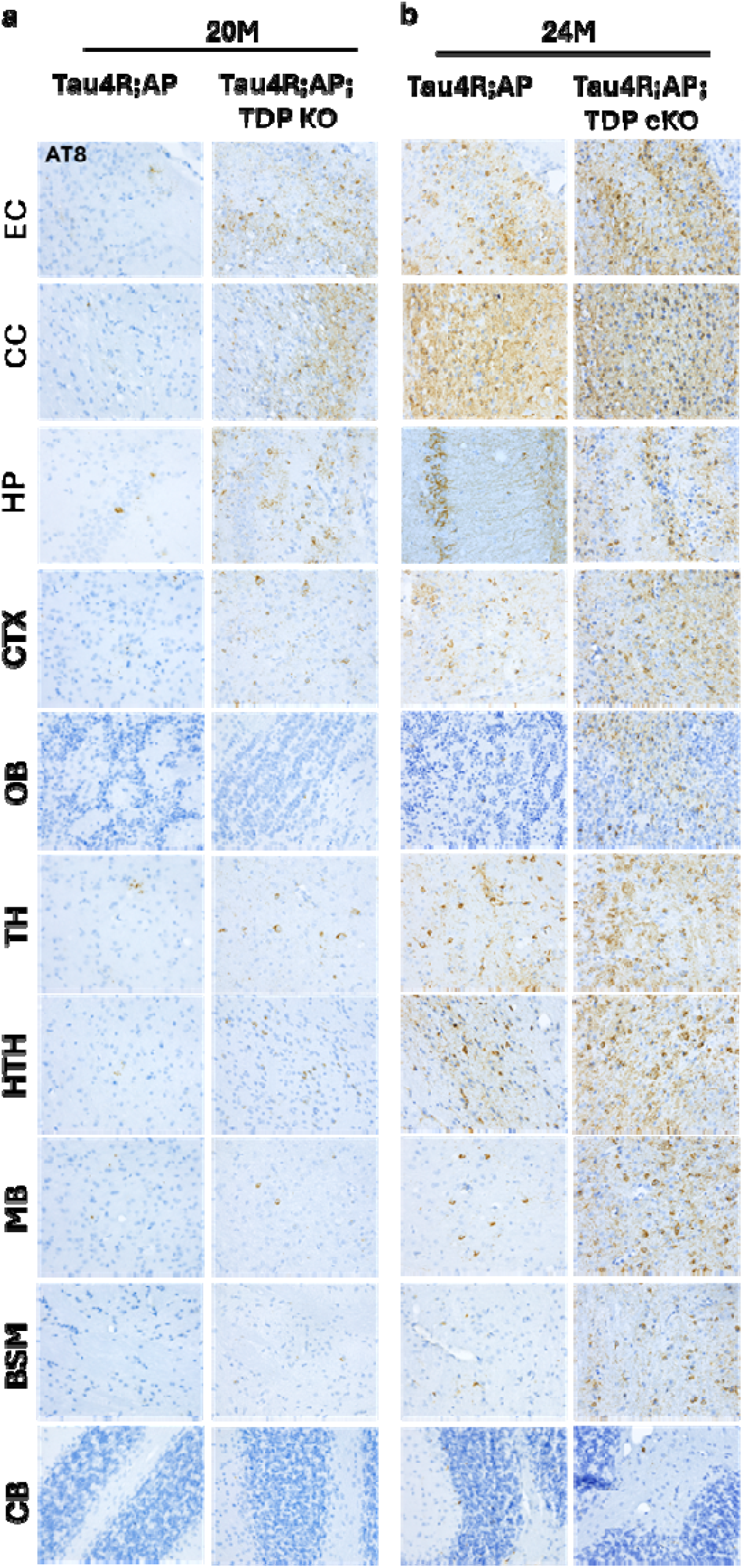
Loss of TDP-43 function accelerates pathological conversion and spread in age dependent manner in the brain of *Tau4R;AP;TDP cKO* mice: (**a-b**) Immunohistochemical analysis extended panel of brains of 20 and 24-month-old Tau4R;AP and Tau4R;AP;TDP cKO using specific antibody AT8 to show the accelerated pathological conversion of tau and spread in brain regions (EC, CC, HP, CTX, OB, TH, HTH, MB, BSM and CB) in age dependent manner.

**Sup Fig. 5:**
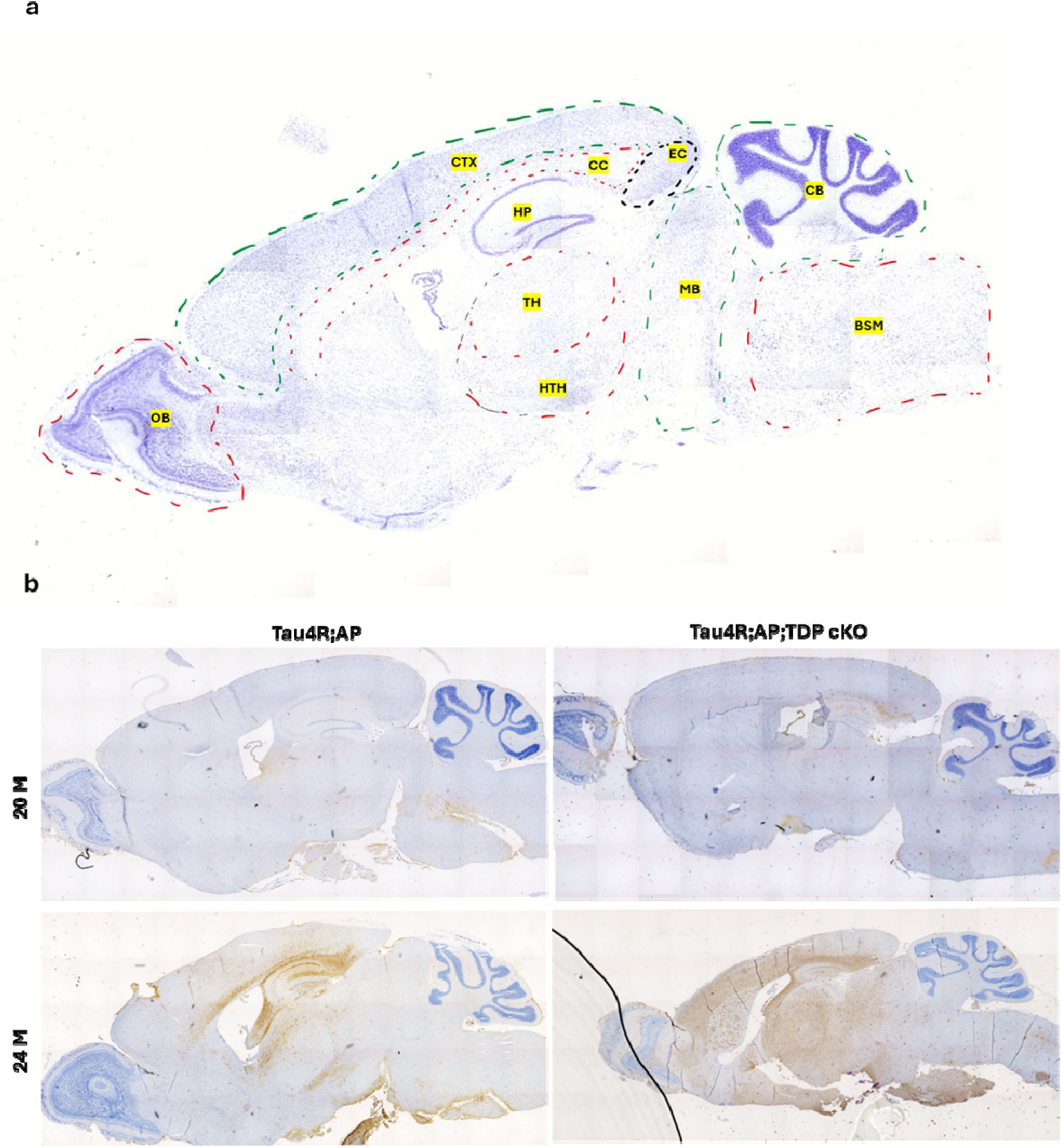
Loss of TDP-43 function accelerates pathological conversion and spread in age dependent manner in the brain of *Tau4R;AP;TDP cKO* mice: **(a)** Nissl staining stained brain section depicted with annotations of different brain regions (EC, CC, HP, CTX, OB, TH, HTH, MB, BSM and CB) used for presenting spread of tau and TDP-43 pathology in the brains. **(b**) Immunohistochemical analysis of brains of 20 and 24-month-old Tau4R;AP and Tau4R;AP;TDP cKO using specific antibody AT8 to show the accelerated pathological conversion of tau and spread in brain regions in age dependent manner.

**Sup Fig. 6:**
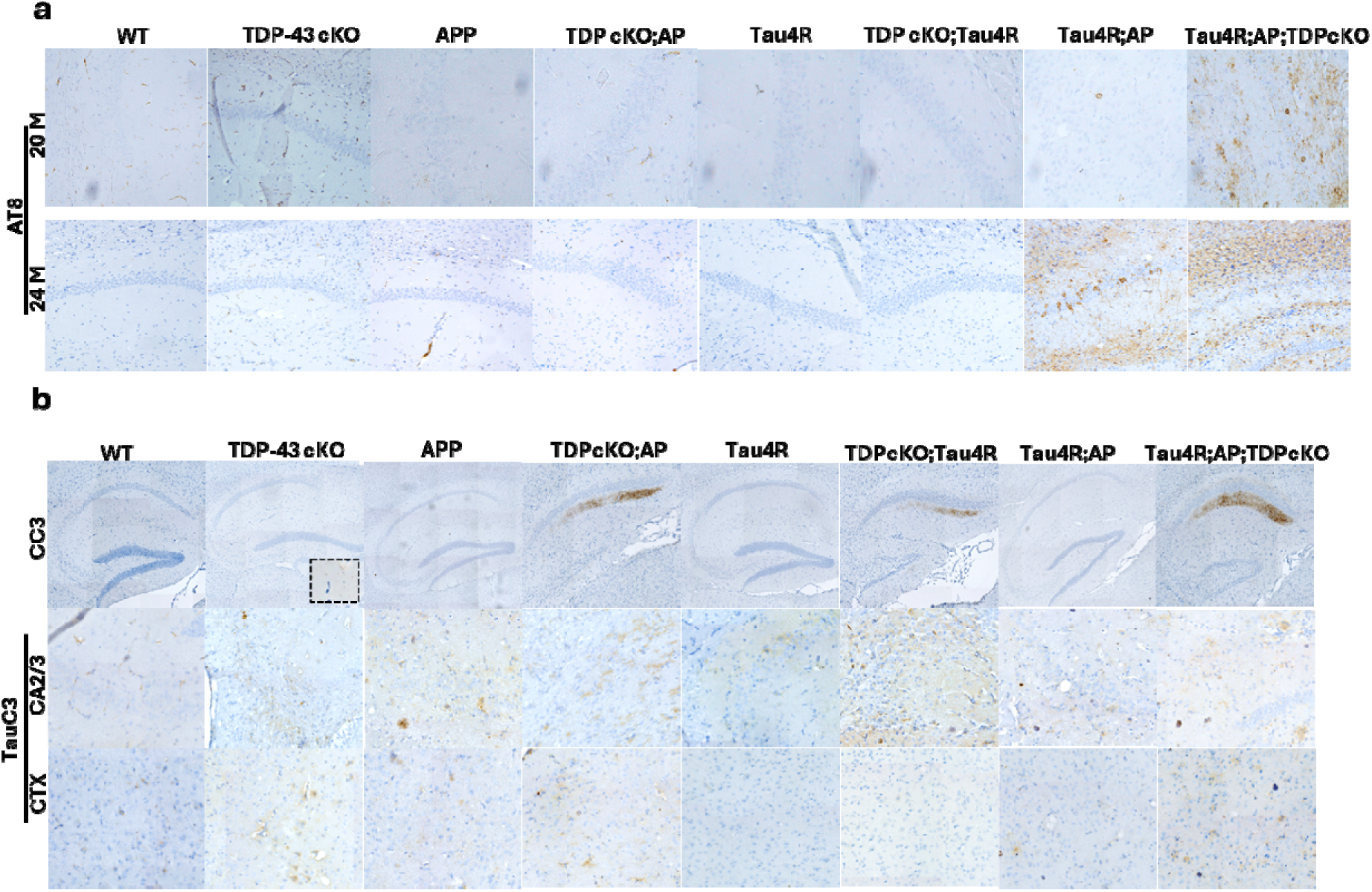
Loss of TDP-43 function accelerates pathological conversion in age dependent manner in the brain of *Tau4R;AP;TDP cKO* mice: **(a)** Immunohistochemical analysis of brains of 20 and 24-month-old WT, APP, TDP-43 cKO, TDP cKO;AP, Tau4R, TDP cKO;Tau4R, Tau4R;AP and Tau4R;AP;TDP cKO using specific antibody AT8 to detect pathological tau conversion. **(b)** Immunohistochemical analysis using cleaved caspase 3 (CC3) brain sections of WT, APP, TDP-43 cKO, TDP cKO;AP, Tau4R, TDP cKO;Tau4R, Tau4R;AP and Tau4R;AP;TDP cKO 20-month-old mice (upper row). The immunoreactivity of cleaved tau (TauC3), in hippocampus and cortex of extended panels of different genotype mice.

**Sup Fig. 7:**
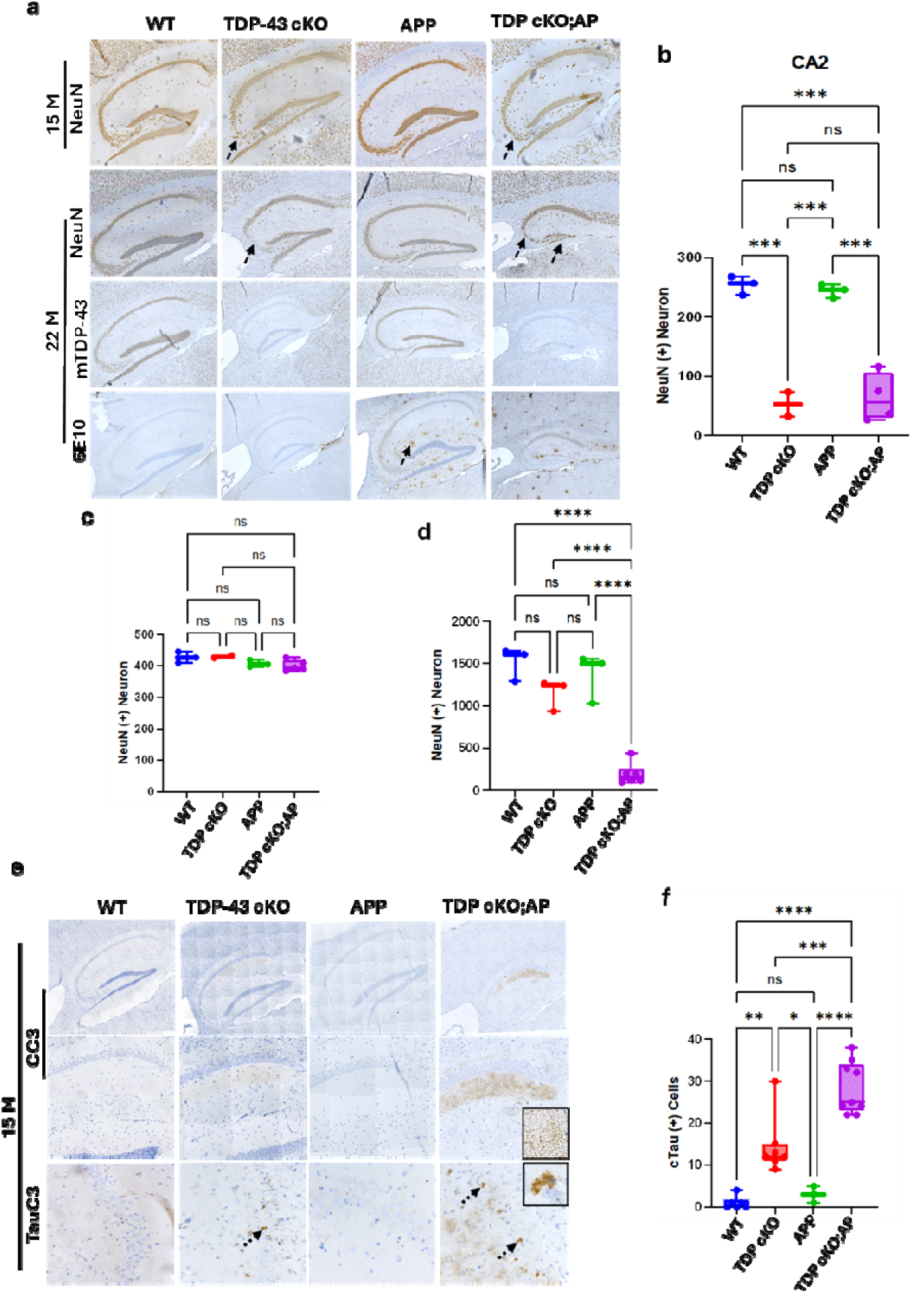
Loss of TDP-43 function accelerates caspase activation-mediate tau cleavage and exacerbate neuron loss in TDP cKO;AP mice: (**a**) Immunohistochemical analysis of brains of 15 and 22-month-old *WT* (n=3), TDP-43 cKO (n=3), AAP (n=3) and *TDP cKO;AP* mice (n=5) using antibody specific to NeuN to detect neurons (Arrow heads indicates neuronal loss in hippocampal subfields). Immunohistochemistry of mTDP-43 and amyloid-β (6E10) showing specific loss of TDP-43 (third row), and presence of amyloid-β deposition in brain of APP and TDP cKO;AP mice (fourth row, arrow heads indicate the amyloid-β plaques). (**b-d**) Quantification of NeuN positive neurons from CA2/3, CA1 and DG respectively in 22-month-old mice (one-way ANOVA; ns: no significant difference; *P<0.05; ****P<0.0001). (**e**) Immunohistochemistry of cleaved caspase 3, in low (first row) and high power (middle row) and cleaved tau (TauC3) in lower row (Arrow heads indicate the immunoreactivity signal of TauC3). (**f**) Quantification of TauC3 positive cells in the hippocampus of 22-month-old mice (one-way ANOVA; ns: no significant difference; *P<0.05; ****P<0.0001).

**Sup Fig. 8:**
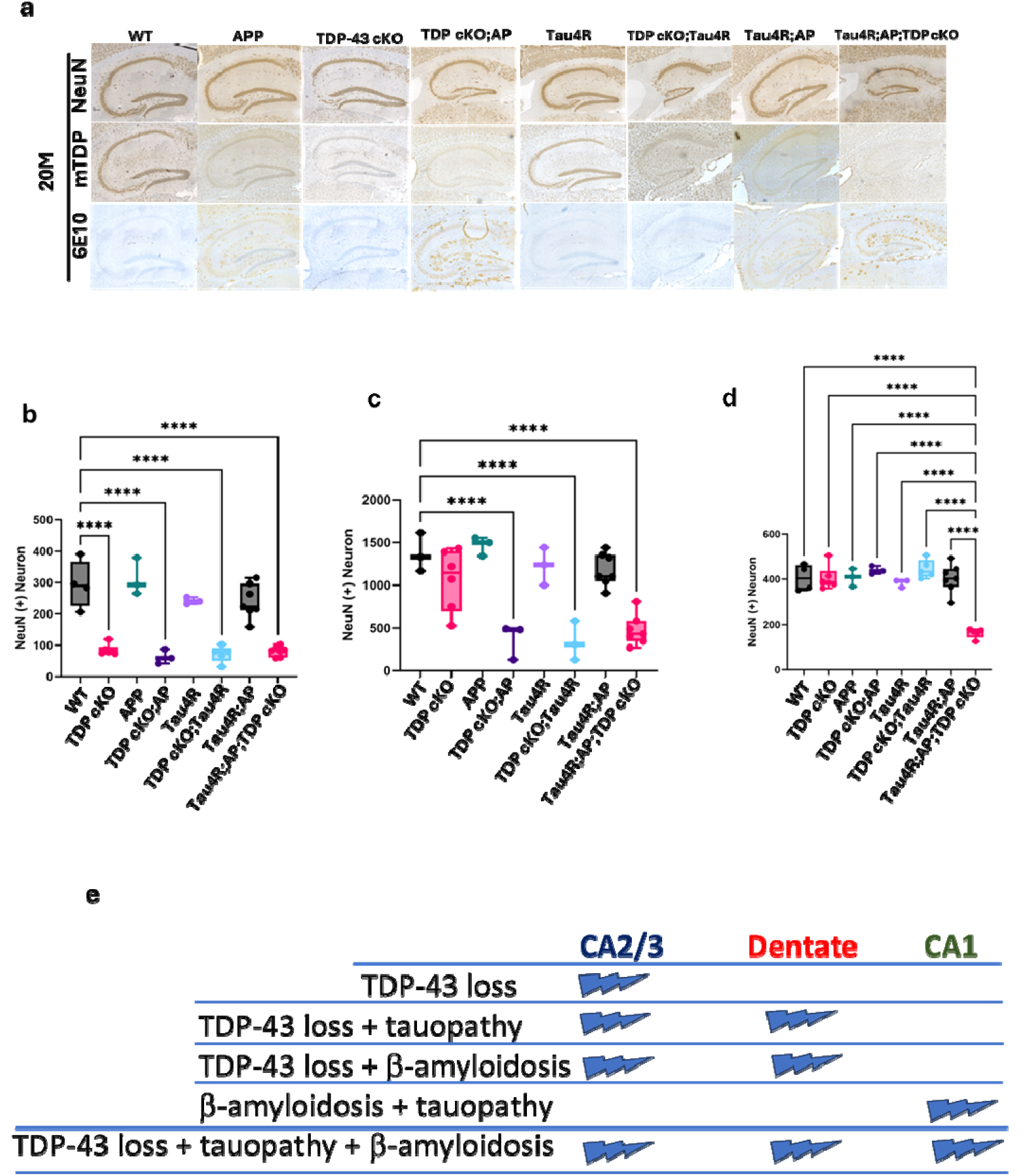
Loss of TDP-43 function exacerbates neuron loss in Tau4R;AP;TDP cKO mice: **(a)** Immunohistochemical analysis of brains of 20-month-old WT (n=4), APP (n=3), TDP-43 cKO (n=4), TDP cKO;AP (n=3), Tau4R (n=3), TDP cKO;Tau4R (n=3), Tau4R;AP (n=5) and Tau4R;AP;TDP cKO (n=4) using antibody specific to NeuN to detect neurons (first row), mTDP-43 for TDP-43 depletion (middle row) and 6E10 for amyloid-β deposition (lower row). **(b-d)** Quantification of NeuN positive neurons in CA2/3, DG and CA1 respectively of 20-months-old mice. **(e)** Graphical representation showing the specific neuronal population in the hippocampus subfield is affected by specific pathology. Hippocampal CA2/3 neurons are selective vulnerable to loss of TDP-43 function, which extends the loss of dentate granular neurons with co-pathology of amyloid and tau. As observed, hippocampal CA1 neurons are selective vulnerable to amyloid and tau pathology, in the all three pathologies tau, amyloid with loss of TDP-43 show mice show broadened degeneration in CA2/3, DG including CA1.

**Sup Fig. 9:**
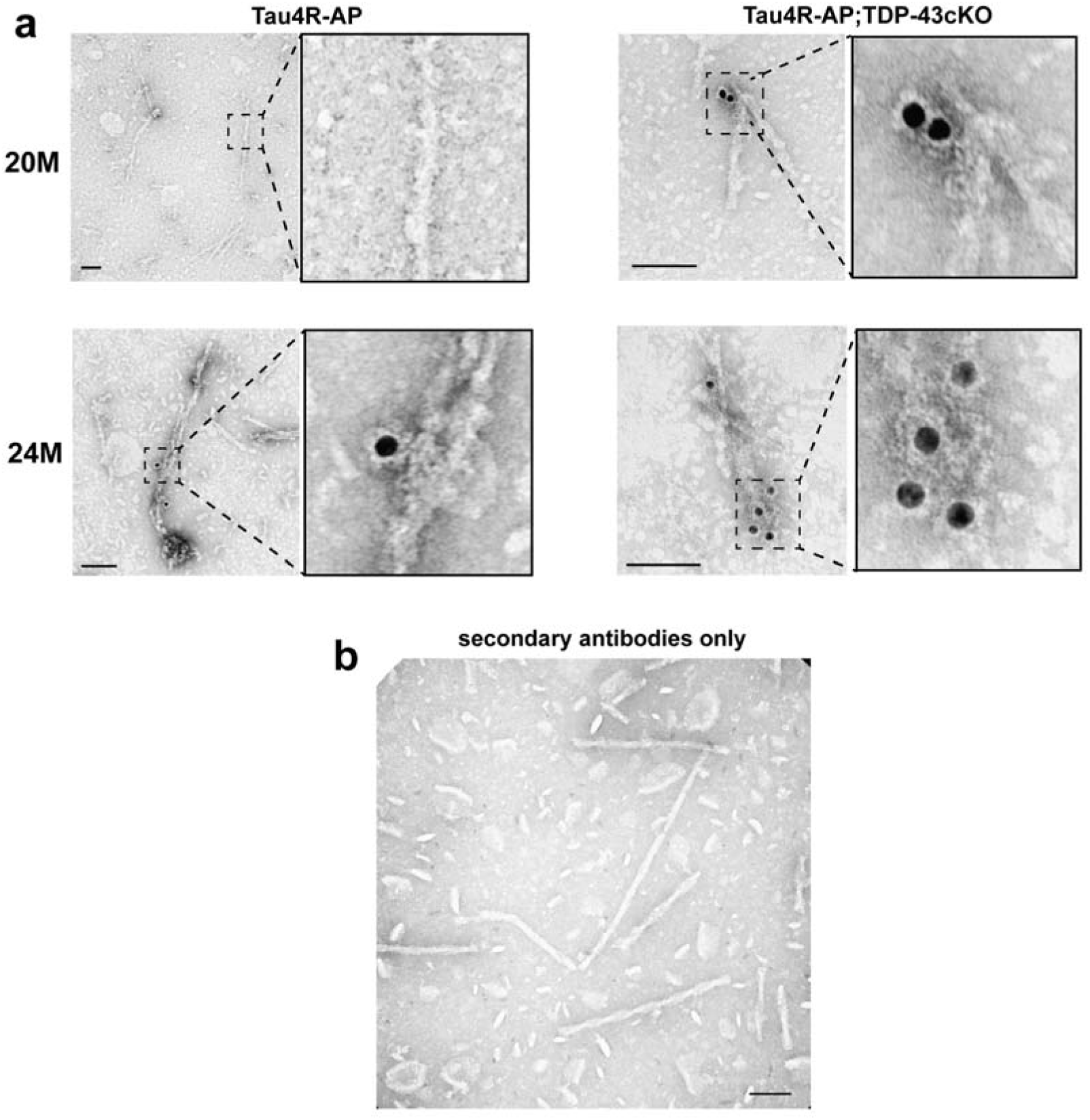
pTDP-43 labeled filaments in Tau4R-AP and Tau4R-AP;TDP-43cKO mice. **(a)** Immuno-EM analysis of sarkosyl-insoluble brain extracts derived from 20 and 24-month-old Tau4R-AP and Tau4R-AP;TDP-43cKO whole brain samples using a primary antibody against TDP-43 (pS409/410). (b) Double labeling immuno-EM analysis of an AD-LATE amygdala sample using goat anti-rabbit 6nm gold-conjugated and goat anti-mouse 12nm gold-conjugated secondary antibodies only to demonstrate the specificities of the tau (pS422) and TDP-43 (pS409/410) primary antibodies (scale bar, 100nm).

**Sup Fig. 10:**
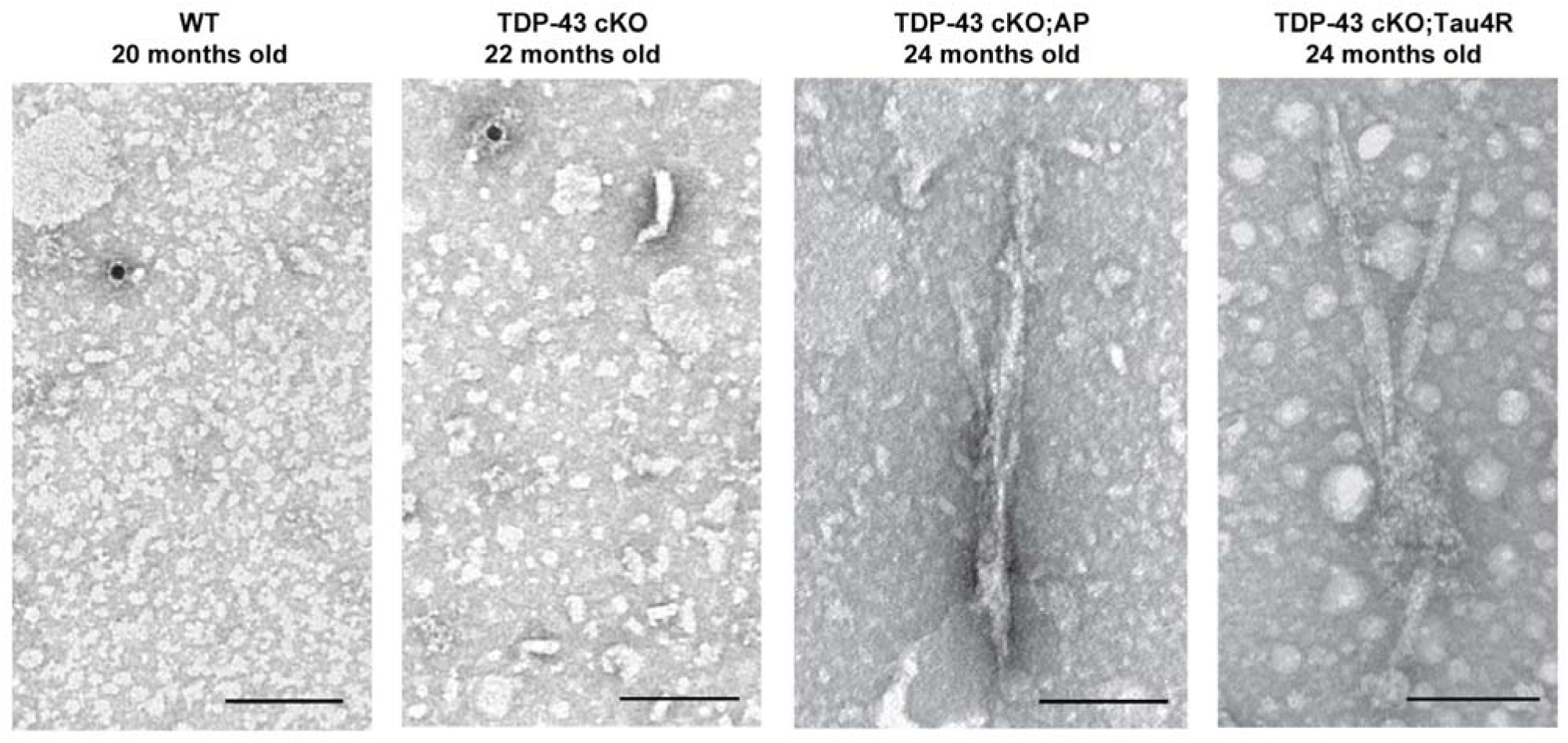
Tau and TDP-43 filaments are absent in control mice. Double labeling immuno-EM analysis of wild-type, TDP-43 cKO, TDP-43 cKO;AP and TDP-43 cKO;Tau4R mouse whole brain samples using primary antibodies against tau (pS422) and TDP-43 (pS409/410) and goat anti-mouse 6nm gold-conjugated and goat anti-rabbit 12nm gold-conjugated secondary antibodies (scale bar, 100nm).

**Sup Fig. 11:**
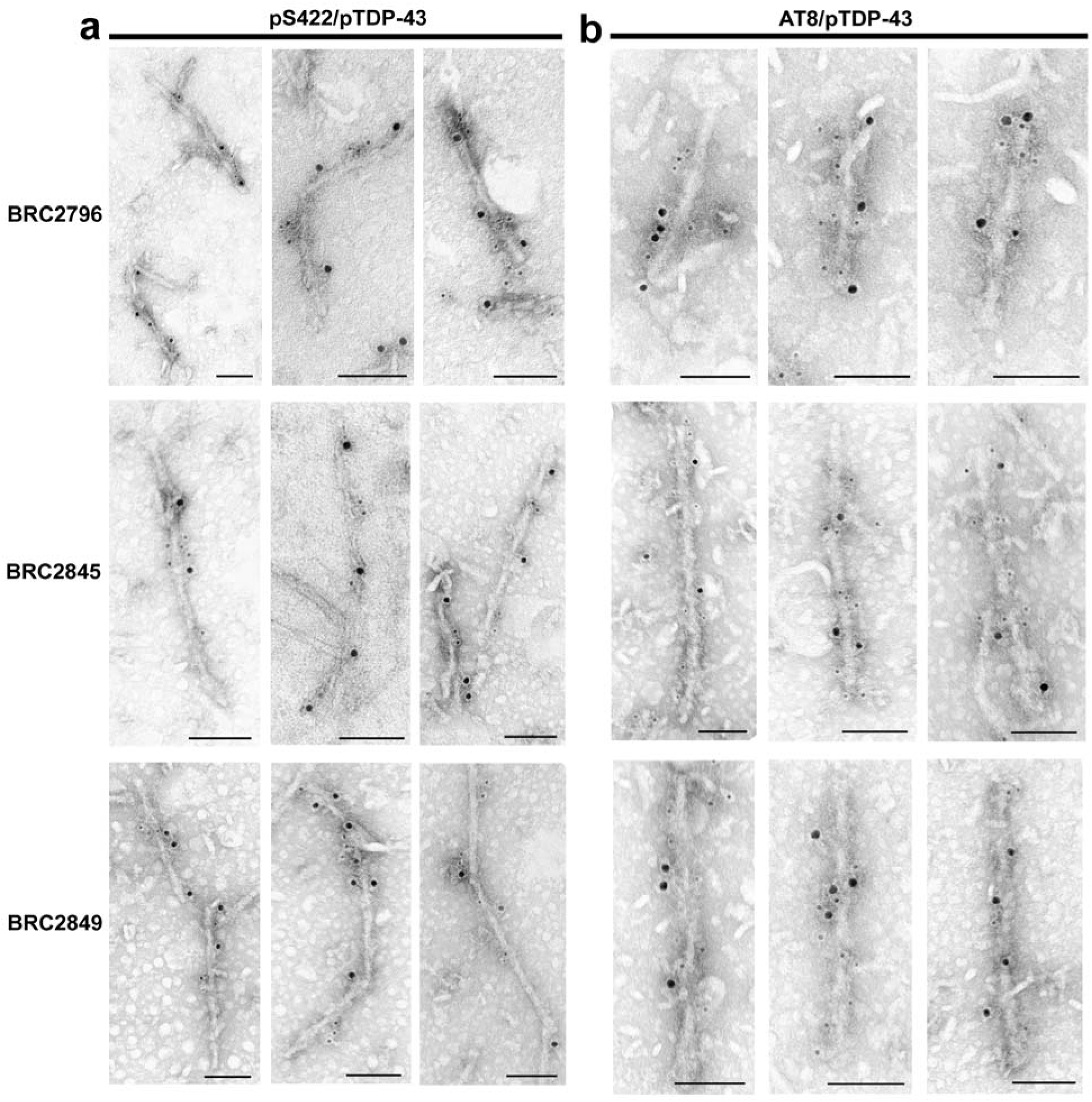
Tau and TDP-43 co-filaments in three additional AD-LATE cases. Double labeling immuno-EM analysis of AD-LATE amygdala samples using primary antibodies against **(a)** tau (pS422) and TDP-43 (pS409/410) or **(b)** AT8 and TDP-43 (pS409/410) and goat anti-mouse 6nm gold-conjugated and goat anti-rabbit 12nm gold-conjugated secondary antibodies (scale bar, 100nm).

**Sup Fig. 12:**
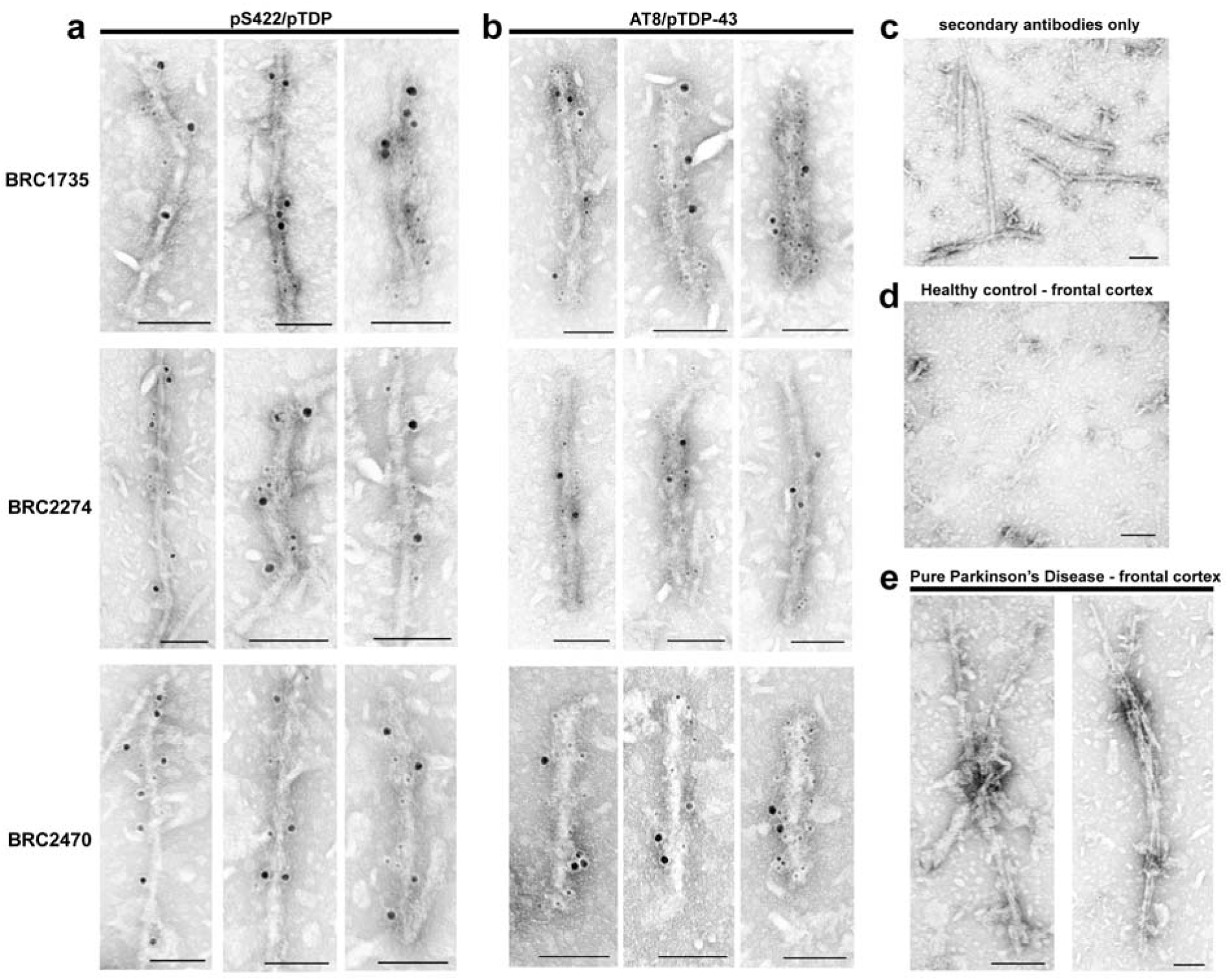
Tau and TDP-43 co-filaments in three additional AD cases. Double labeling immuno-EM analysis of AD amygdala samples using primary antibodies against **(a)** tau (pS422) and TDP-43 (pS409/410) or **(b)** AT8 and TDP-43 (pS409/410) and goat anti-mouse 6nm gold-conjugated and goat anti-rabbit 12nm gold-conjugated secondary antibodies. **(c)** Double labeling immuno-EM analysis of AD amygdala sample using goat anti-mouse 6nm gold-conjugated and goat anti-rabbit 12nm gold-conjugated secondary antibodies only. (**d-e**) Double labeling immuno-EM analysis of frontal cortex samples from healthy control and pure Parkinson’s disease cases using primary antibodies against tau (pS422) and TDP-43 (pS409/410) and goat anti-mouse 6nm gold-conjugated and goat anti-rabbit 12nm gold-conjugated secondary antibodies (scale bar, 100nm).

